# Dissecting pOXA-48 fitness effects in clinical enterobacteria using plasmid-wide CRISPRi screens

**DOI:** 10.1101/2025.01.23.633114

**Authors:** Alicia Calvo-Villamañán, Jorge Sastre-Dominguez, Álvaro Barrera-Martín, Coloma Costas, Álvaro San Millan

**Affiliations:** Centro Nacional de Biotecnología (CNB-CSIC), Madrid, Spain; Centro de Investigación Biológica en Red de Epidemiología y Salud Pública, Instituto de Salud Carlos III, Madrid, Spain

## Abstract

Conjugative plasmids are the main vehicle for the spread of antimicrobial resistance (AMR) genes in clinical bacteria. AMR plasmids allow bacteria to survive antibiotic treatments, but they also produce physiological alterations in their hosts that commonly translate into fitness costs. Despite the key role of plasmid-associated fitness effects in AMR evolution, their origin and molecular bases remain poorly understood. In this study, we introduce plasmid-wide CRISPR interference (CRISPRi) screens as a tool to dissect plasmid-associated fitness effects. We designed and performed CRISPRi screens targeting the globally distributed carbapenem resistance plasmid pOXA-48 in 13 different multidrug resistant clinical enterobacteria. Our results revealed that pOXA-48 gene-level effects are conserved across clinical strains, and exposed the key role of the carbapenemase-encoding gene, *bla*_OXA-48_, as the main responsible for pOXA-48 fitness costs. Moreover, our results highlighted the relevance of postsegregational killing systems in pOXA-48 vertical transmission, and uncovered new genes implicated in pOXA-48 stability. This study sheds new light on the biology and evolution of carbapenem resistant enterobacteria and endorses CRISPRi screens as a powerful method for studying plasmid-mediated AMR.

## Introduction

Antimicrobial Resistance (AMR) is quickly becoming one of the most concerning global health problems, with 1.2M deaths worldwide associated with AMR infections each year^1^. Studies have estimated that in a few decades AMR infections will be the main cause of non-natural death, increasing the yearly toll to 10M deaths associated with AMR^1,2^ and causing an annual loss of 1% of the world GDP^3^. AMR is particularly alarming in the clinical setting due to the generalised use of antibiotics in hospitals, which expedites the evolution of antimicrobial resistant bacteria^4^. However, the vast majority of scientific studies on AMR evolution have been performed with laboratory bacterial strains, which don’t fully represent the complexity of clinical ones. We therefore are in dire need of more scientific studies performed in clinical strains, ones that are able to capture AMR in its full, real-world complexity^5^.

AMR can be achieved through different mechanisms^6^. Out of them, conjugative plasmids play an essential role in the spread of AMR among clinical enterobacteria^7^. These extrachromosomal genetic elements often carry AMR genes and have the ability to be transferred horizontally to cells that are nearby, promoting the dissemination of AMR genes across bacterial populations^8^. Plasmids can be double-edged swords for the bacteria that carry them. On the one hand, they carry genes that can be useful in the right conditions *(e.g.* a specific AMR gene when a given antibiotic is prescribed to a patient). On the other hand, they alter the fine-tuned metabolism of the cell, often causing changes to the fitness of their bacterial hosts^9^. Multiple studies have shown plasmids to produce variable fitness effects across bacterial hosts^7,10–12^. These host-specific responses to carrying the same plasmid help explain why plasmids are frequently associated with a small fraction of hosts, regardless of their often wide host range.

Plasmid-associated fitness costs arise from genetic conflicts between the novel plasmid genes and the genetic background of the host^13,14^. The molecular mechanisms behind these interactions are multifactorial and poorly understood^15,16^. Plasmid’s costs can originate during the different phases of their life-cycle in the cell: from plasmid reception^17^, to replication^18^, gene expression^19^ and conjugation^20^.

However, recent studies suggest that the main source of plasmid fitness costs are the interactions between plasmid-encoded proteins and the host bacterium. For example, plasmids often encode for homologues of bacterial regulators which they can use to re-tune the regulatory networks of the bacterium to their advantage^16,21^. While beneficial to plasmid survival, these changes in metabolic flux might affect essential cellular processes and create a fitness cost for the host. Similarly, by encoding extra copies of dose-sensitive bacterial genes, plasmids might disturb the optimal protein concentration, leading to a fitness cost^19^. Finally, by encoding homologues of highly connected bacterial proteins, multiple metabolic processes might be affected at once in the host^22^. Several studies have centred on trying to understand how plasmid fitness costs play a role in plasmid evolution. A trade-off between horizontal and vertical transmission of the plasmid has been proposed: a very efficient plasmid conjugation machinery tends to be costly to the plasmid host^10,23^. A second trade-off between the expression of resistance genes and vertical transmission has also been proposed: in the absence of antibiotics the costs associated with plasmids have been shown to often arise from the expression of their resistance genes^24–26^.

pOXA-48 is a highly-conjugative plasmid of great clinical relevance, which promotes the dissemination of the carbapenemase-encoding gene *bla*_OXA-48_ across clinical enterobacteria^27,28^. Carbapenemases are able to hydrolyse carbapenems, which are used in hospitals as last-resort treatments of multidrug resistant (MDR) infections. Because of this, pOXA-48 dissemination in the nosocomial context represents a global public-health threat. Despite its family-wide host range, clinical data shows the plasmid to be strongly associated with specific high-risk *Klebsiella pneumoniae* clones (such as those belonging to sequence type ST11^29,30^).

In this work we use plasmid-wide CRISPR interference (CRISPRi) screens to investigate the pOXA-48 gene-level fitness effects in clinical strains of enterobacteria. Our results show that although carrying pOXA-48 leads to very different fitness responses in its different hosts, the gene-level effects are conserved amongst them. We confirm the importance of the toxin-antitoxin (TA) system PemI-PemK in the vertical transmission of the plasmid, and highlight the previously unknown role of plasmid-encoded modulators of expression in pOXA-48 stability in *E. coli*. Our data also suggests that pOXA-48 might be better adapted to *K. pneumoniae* than to *Escherichia coli,* which could explain its strong association in the clinic with the first species. Most importantly, we show that the carbapenemase OXA-48 is the main responsible for the fitness costs associated with the plasmid.

## Results

### Designing a pOXA-48-wide CRISPRi screen for clinical enterobacteria

We started by designing and building a CRISPRi set-up to study gene-level fitness effects of pOXA-48 in clinical enterobacteria. CRISPRi works by using dCas9, a DNA-binding protein that can be directed to virtually any region of DNA guided by a small RNA molecule named single-guide RNA (sgRNA). Once bound to DNA, dCas9 inhibits the expression of the targeted region (Fig. 1A). In CRISPRi screens, a pool of bacteria is built so that each individual bacterium carries dCas9 programmed to silence an individual gene through a specific sgRNA. This pool of bacteria can then be propagated, and the fitness effects of each gene in the given context can be determined through the change in frequency of gene-specific sgRNAs in the population (Fig. 1B). For the pOXA-48-wide CRISPRi screen, we designed 5 sgRNAs targeting each plasmid gene and up to 10 sgRNAs per intergenic region (depending on the size of the region, S1 Table). Each sgRNA was cloned in pFR56apm, a vector expressing the CRISPRi machinery under DAPG induction. The resulting 568-plasmid library was named OXAlib.

**Fig. 1.**
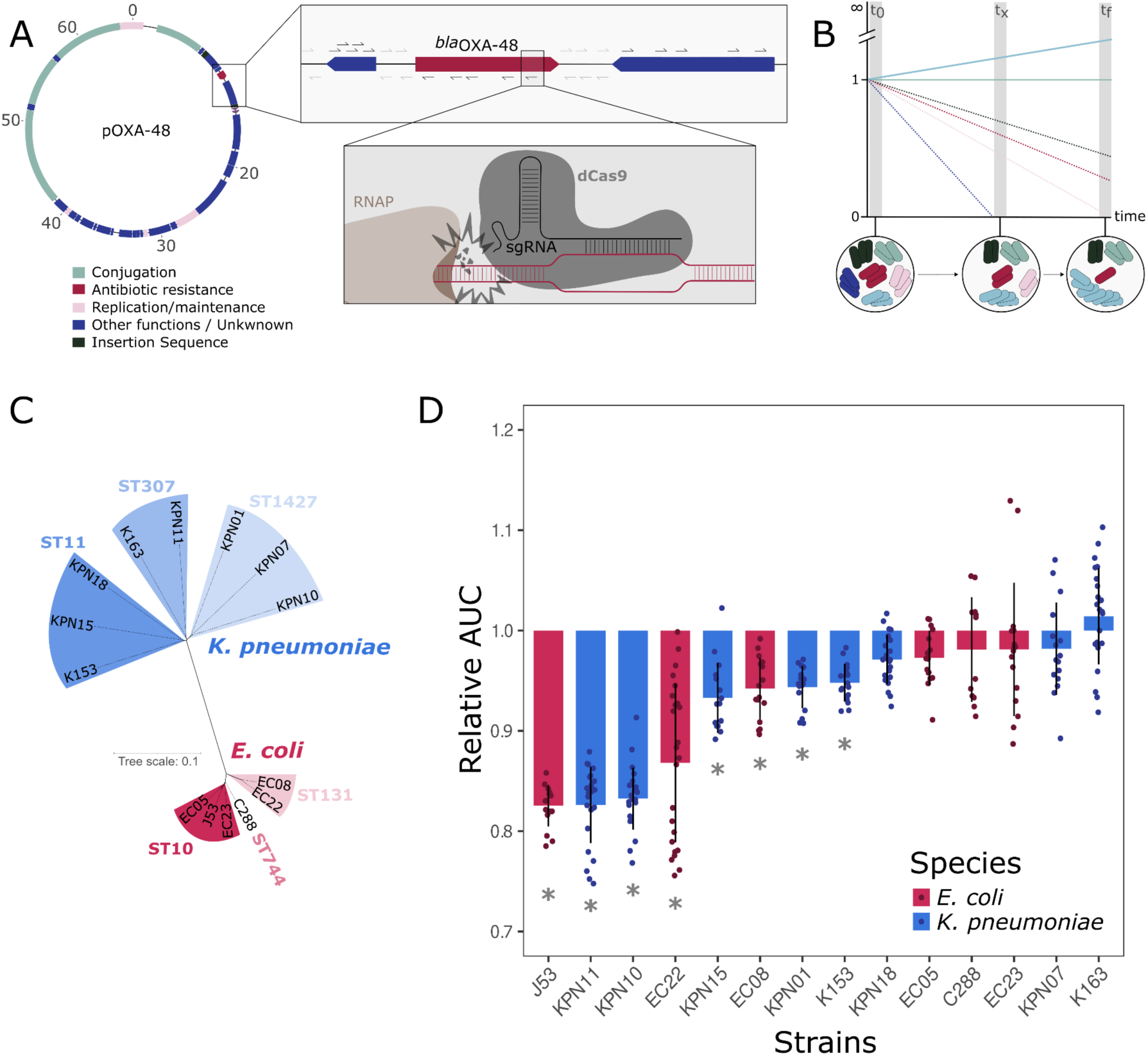
CRISPRi screens set-up and fitness effects of pOXA-48. Depiction of the mechanism of CRISPRi screens and fitness costs associated with carrying pOXA-48 in the strains under study. (A) Schematic representation of CRISPRi gene-silencing and the set-up of CRISPRi screens. Values in pOXA-48 correspond to Kb. Each individual arrow represents an sgRNA targeting the region, and the orientation of the arrow indicates the orientation of dCas9 when binding to the DNA strand (RNAP stands for RNA polymerase). (B) Schematic representation of CRISPRi screens. Each colour corresponds to bacteria with a different pOXA-48 gene silenced (*i.e.* carries pFR56apm with an sgRNA programmed against said gene). Bacteria which carry a guide silencing a beneficial gene will decrease in frequency the population during the screening, lowering the amount of the guide over time (e.g. red cells). On the other hand, bacteria carrying guides which silence costly genes will be enriched during the screening (e.g. light blue cells) (C) Unrooted phylogenetic tree of the enterobacteria clinical strains used in this study. Multi Locus Sequence Types (STs) are indicated. (D) Distribution of pOXA-48 fitness effects (Relative Area Under the Growth Curve [see Methods]) in the clinical strains selected for the CRISPRi screening. Asterisks indicate significant cost of pOXA-48 in each of the strains (paired t-tests against each relative plasmid-free strain after Bonferroni correction; n = 16; p < 0.05).

We selected a collection of 13 clinical strains of *K. pneumoniae* (n=8) and *E. coli* (n=5), representative of the diversity of MDR enterobacteria isolated from hospitalised patients at a large hospital in Madrid^30^, to perform the CRISPRi screen. This collection included clones that are typically associated with pOXA-48 in our hospital and others, such as *K. pneumoniae* ST11, ST307 and *E. coli* ST10^30^ (Fig. 1C). As a control we also included *E. coli* J53, a lab strain derived from K12^31^. For more information on the strains used, consult the supplementary data (S2 Table).

We first determined the fitness effects of pOXA-48 (variant K8, GenBank MT441554) in each individual host, by comparing the growth of pOXA-48 carrying and pOXA-48-free isogenic versions of each strain (Fig. 1D). As previously described, the plasmid produced variable fitness effects in the 14 strains^12,32^. Prior to performing the CRISPRi screens, we confirmed the efficiency of dCas9-mediated silencing in all strains, by targeting the essential gene *rpsL*^33^ with a guide expressed from the pFR56apm vector (SFig. 1).

### CRISPRi screens reveal conserved pOXA-48 gene-level effects across clinical strains

We introduced the OXAlib library in the 14 pOXA-48-carrying strains and performed the CRISPRi screens with continuous dCas9 expression for ∼85 generations (72 h). The screens were performed in triplicate, both without antibiotic pressure and in the presence of sub-inhibitory concentrations of ertapenem, to which pOXA-48 confers resistance. Due to the differences in MIC for the different strains, different concentrations of ertapenem were used for each strain (S1 Table). To determine the fitness effects of pOXA-48-encoded genes, we collected samples at 0, ∼25, ∼55 and ∼85 generations and measured the log2 fold-change (log2FC) of sgRNAs in the population. The change in frequency of gene-specific sgRNAs in the population informed the fitness effects associated with silencing each particular gene. The data shown in this paper corresponds to ∼85 generations, as that’s when the effect of each sgRNA was most visible. In parallel, the stability in the population of both pOXA-48 and pFR56apm were also assessed (SFig. 2).

Despite the variable fitness effects produced by pOXA-48 across the clinical strains (Fig. 1D), we observed that silencing each pOXA-48 gene individually led to similar fitness effects across strains (Fig. 2A). To confirm this observation, we correlated the log2FC of all the genes and intergenic regions between all the strains per experimental condition. Pearson correlation indexes (*r*) resulted in highly positive values for most of the pairwise comparisons between strains both in absence and presence of ertapenem (0.25 < *r* <0.90, median = 0.64 without ertapenem; −0.07 < *r* < 0.91, median = 0.66 with ertapenem. Fig. 2B; S3 Table).

**Fig. 2.**
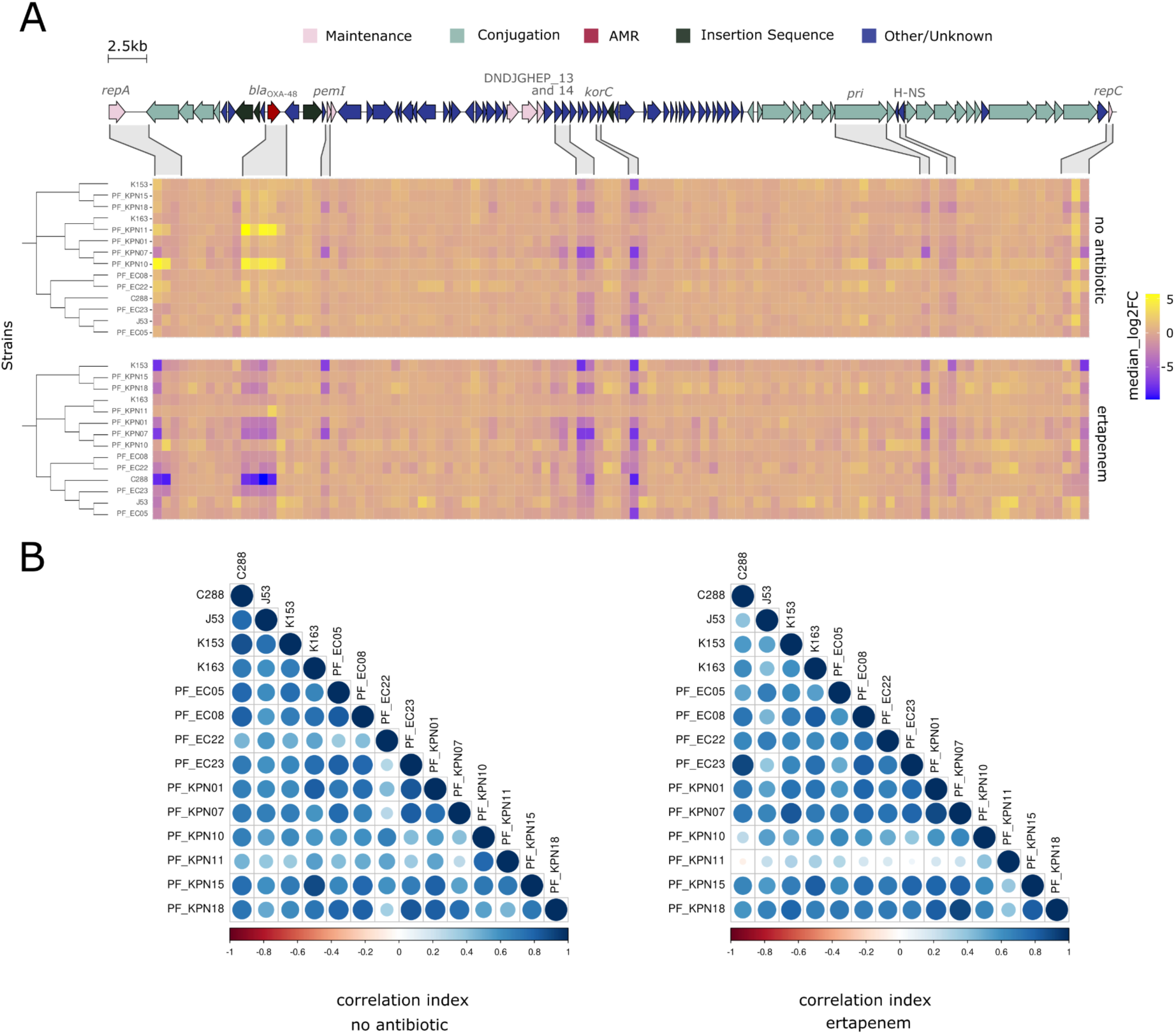
pOXA-48-encoded genes produce similar fitness effects across clinical strains. Representation of the fitness effects associated with silencing each gene in pOXA-48 individually in the presence and absence of antibiotic pressure, and how conserved those effects are amongst strains. (A) Heatmap of CRISPRi gene scores at the end of the screening (t = 72 h, ∼85 generations). Fitness effects associated with silencing each individual gene / intergenic region in the absence of selection (top panel) and in the presence of ertapenem (bottom panel). The score for each gene / intergenic region corresponds to the median log_2_ fold-change of the sgRNAs targeting the element in the population (see Methods). Blue shades indicate those genes which silencing is detrimental (i.e. genes that are beneficial in that condition), whereas yellow shades correspond to genes which silencing is beneficial during the experiment (i.e. genes that are costly in that condition). Coding regions of pOXA-48 are shown with different colours indicating common functional clusters as shown on top. Genes and surrounding intergenic regions showing a significant deviation in the CRISPRi screening score in both conditions are highlighted in the top. A fully annotated version of the figure is available in SFig. 4. (B) Pearson correlation matrices between strains’ gene scores at the end of the screening without (left) and with ertapenem (right). Blue shades indicate positive Pearson correlation coefficients, meaning similar tendencies in the overall screening results between the strains. Circle sizes indicate correlation absolute values from 0 to 1 (blue) or −1 to 0 (red).

Finally, we aimed to confirm that the overall gene-level effects revealed in CRISPRi screens were able to recapitulate the fitness effects produced by the entire pOXA-48 plasmid. Our results showed that the median log2FC of plasmid-targeting sgRNAs in the absence of ertapenem (compared to the control guides) anticorrelated with the relative AUC of pOXA-48-carrying strains (*R* = −0.73, p-value = 0.0045). Therefore, silencing plasmid genes is more beneficial as the plasmid increases in cost. These results confirmed that our CRISPRi data is able to reflect overall plasmid fitness effects (SFig. 3).

### Analysing pOXA-48 gene-level fitness effects

To identify which genes play a significant role in pOXA-48 fitness effects, we performed a permutation test to pinpoint the genes whose silencing led to statistically significant changes in frequency both in presence and absence of ertapenem (Fig. 3A). These genes were (i) *bla*_OXA-48_, the β-lactamase gene; (ii) *repA*, the replication initiation gene (as representative of the *repCBA* locus); (iii) *pri,* which is involved in conjugation, and shows homology to DNA primases; (iv) *korC*, a transcriptional repressor with unknown function in pOXA-48; (v) a H-NS-like family regulator gene; (vi) an operon with unknown function formed by DNDJGHEP_0013 and _0014, predicted to be an ABC-transporter substrate binding protein and an Xre-like transcriptional regulator, respectively; and (vii) *pemI*, the antitoxin of the *pemI-pemK* system, responsible for post-segregational killing in case of plasmid mis-segregation^34^. In line with our previous results (Fig. 2), the genes showing significant effects were highly conserved across strains (SFig. 5). Interestingly, the response was more generally conserved in *K. pneumoniae* than in *E. coli* (Welch two sample t-test, t = −2.1694 p-value = 0.04234 without ertapenem; two-sided Kruskal-Wallis test, chi-squared = 5.6326, p-value = 0.01763 with ertapenem; SFig. 5).

**Fig. 3.**
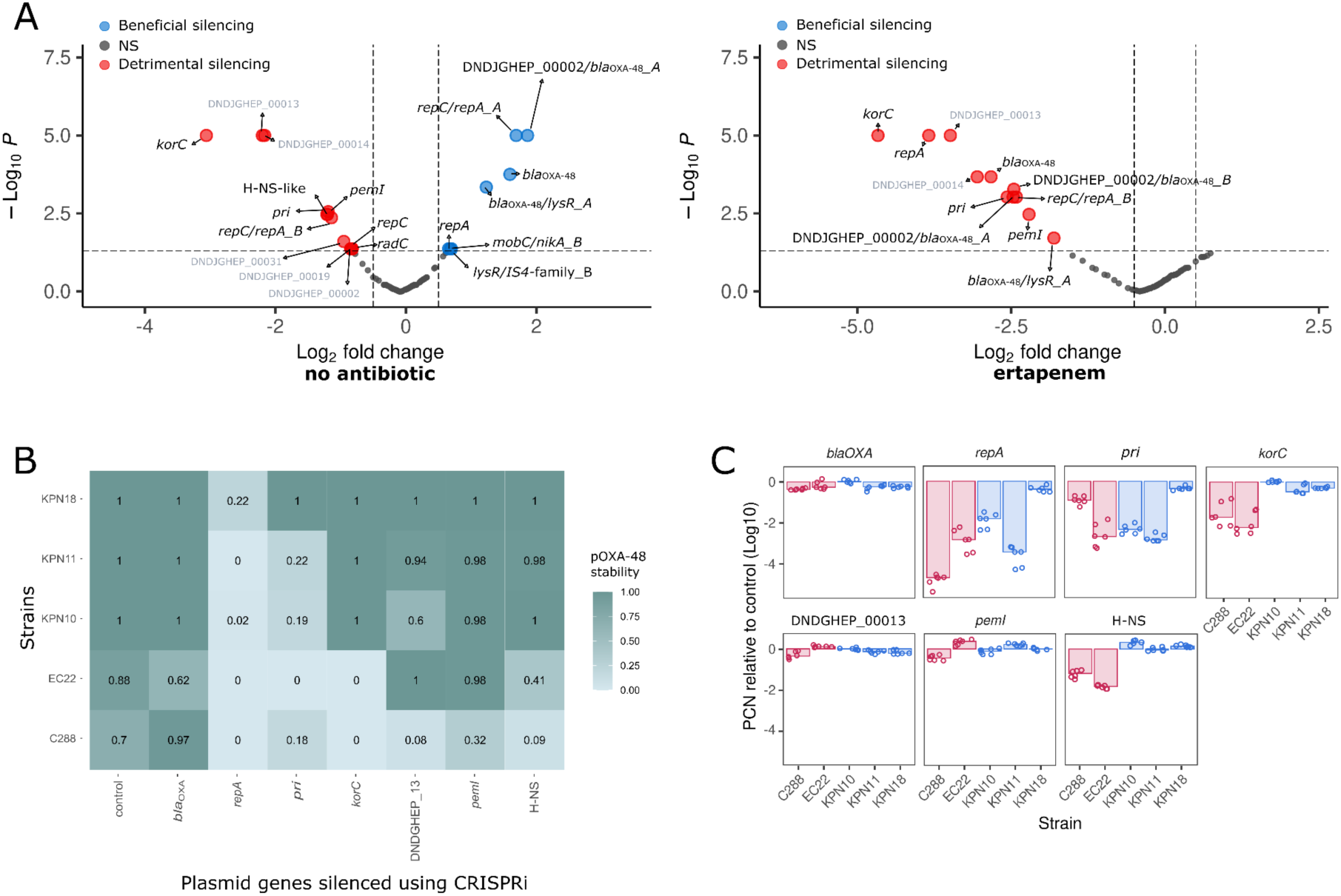
Role of different plasmid genes in pOXA-48 biology. Identification of pOXA-48 genes producing significant fitness effects and their effect on plasmid stability. (A) Volcano plots obtained from permutation test assays in absence of selection (left) and in the presence of ertapenem (right). The Y axis shows the logarithmic transformation of the p value from the permutation test, while the x axis indicates log_2_ fold-change values from CRISPRi screens. Grey points indicate genes and intergenic regions which silencing was non-significant (i.e. −0.5 < log_2_ fold-change < 0.5; and p > 0.05). Genes and intergenic regions which silencing was detrimental in each condition are indicated in red, whereas genes which silencing was beneficial are coloured in blue. The labels of genes encoding hypothetical proteins are shown in grey. (B) pOXA-48 stability after 24 h of CRISPRi gene silencing of individual plasmid genes (X axis) in a subset of 5 strains (Y axis; 2 *E. coli*: EC22 and C288; and 3 *K. pneumoniae*: KPN18, KPN11 and KPN10) from our screenings (in presence of apramycin to select for pFR56apm). (C) pOXA-48 Copy Number measured silencing individual plasmid genes relative to control guide values for each strain tested (Log_10_ scale). Negative values indicate PCN decrease when silencing the indicated gene. Each point shows an individual replicate (n = 6). Bars correspond to median relative PCN values for all the replicates in each of the strains tested. Bars shaded in blue correspond to *K. pneumoniae* strains, and in pink to *E. coli* strains.

Our analysis detected genes that, when silenced, produced a significant advantage or disadvantage for the bacterial host. In the absence of ertapenem, silencing five of the genes led to a negative effect (*pri*, H-NS, DNDJGHEP_0013 and _0014, and *pemI*), while silencing of *bla*_OXA-48_ and *repA* led to a positive fitness effect (Fig. 3A, left panel). In the presence of ertapenem, silencing of all seven genes led to a detrimental effect (Fig. 3A, right panel).

### Investigating the role of individual genes on plasmid stability

Our results showed that most of the genes producing a significant signal when silenced led to a fitness disadvantage. For *bla*_OXA-48_, the fitness disadvantage in the presence of ertapenem can be easily explained because the OXA-48 carbapenemase is responsible for the resistance phenotype. In the case of *pemI*, silencing the antitoxin gene probably leads to a PemK-mediated toxic effect (although this is likely mitigated by the polar silencing of *pemK*, since *pemI-pemK* form an operon^35^). The effect of PemK is analysed in depth in the following section. For the remaining genes, we hypothesised that the fitness defects could be due to a reduction in plasmid stability. Specifically, in the presence of ertapenem, plasmid loss will produce an obvious growth defect, since the cells will no longer produce the OXA-48 carbapenemase. Moreover, plasmid loss should also lead to a PemK-mediated post-segregational killing effect, and this should hold true both in presence and absence of ertapenem. Therefore, we decided to investigate the potential role of silencing these genes in pOXA-48 stability. We individually cloned the strongest sgRNA from the CRISPRi screen for each gene in pFR56apm. Then, we transformed the pFR56apm plasmids individually in a subset of the strains used in our screen, namely *K. pneumoniae* KPN10, KPN11 and KPN18, and *E. coli* C288 and EC22. Using these constructions we investigated how blocking the expression of each gene affected plasmid stability.

We blocked the expression of each gene individually over a 24 h growth cycle and measured the stability of pOXA-48 using replica plating (Fig. 3B). We confirmed the phenotypic results by measuring pOXA-48 copy number in the same populations by qPCR (Fig. 3C). Our results showed that silencing all the genes under study, except *bla*_OXA-48_, led to a reduction in pOXA-48 stability compared to the non-targeting control guide at least in one of the strains (one-tailed z-tests after Bonferroni adjustment, n = 5, p < 0.05). The most dramatic decrease in plasmid stability was mediated by silencing *repA*, which led to the complete curing of the plasmid in 4/5 of our strains. This was to be expected, given the role of RepA as the replication initiation protein of pOXA-48. Surprisingly, *pri* seems to play a similar role in plasmid stability as *repA.* This gene had been described as essential for conjugation in a similar plasmid to pOXA-48 belonging to the plasmid taxonomic unit L/M^36^. However, its potential activity as a DNA primase could play a role in plasmid replication as well.

H-NS, *korC* and the DNDJGHEP_0013 and _0014 genes produced a stronger effect in pOXA-48 stability in *E. coli* than in *K. pneumoniae*. These results are further supported by a parallel study working with a transposon insertion library of pOXA-48 in *E. coli* MG1655, in which Baffert *et al.* showed that KorC and DNDJGHEP_0014 (*orf20*) are essential for plasmid stability^37^. The specific mechanisms by which these genes affect plasmid stability are difficult to assess. H-NS, *korC* and DNDJGHEP_0014 (which forms an operon with DNDJGHEP_0013 and is therefore also silenced by guides targeting this upstream gene) encode for transcriptional regulators. Further work will be needed to clarify the transcriptional effects associated to silencing these genes and how they impact plasmid satility. Interestingly, although silencing these genes produced a negative effect in most *K. pneumoniae* strains in our CRISPRi screens, they affected pOXA-48 stability to a lower extent in this second experiment. However, it is important to remember that the results of silencing individual genes were obtained after a single growth cycle (8-10 generations), while the CRISPRi screens were performed for 85 generations. Finally, and as expected, *pemI* produced a very modest effect on plasmid stability, and *bla*_OXA-48_ produced no effect at all.

### PemK enhances vertical transmission of pOXA-48

We hypothesised that PemK was responsible for the fitness defects associated with pOXA-48 loss in the clinical strains under study^34^. To test this hypothesis, we first cloned *pemK* in an expression vector under the control of the pTet promoter. We introduced this vector in *K. pneumoniae* KPN10 and *E. coli* EC22. Interestingly, even if the expression of *pemK* was not induced, we could not obtain viable transformants for KPN10 carrying the intact vector, suggesting that even a leaky expression of *pemK* is lethal in this strain. We tried to transform the vector in a different *K. pneumoniae* strain, KPN11, with the same result. For *E. coli* EC22, we could transform this vector, probably due to a tighter pTet repression in this species. Our results showed that *pemK* expression led to a dramatic growth defect in this strain (SFig. 6). These results confirm that PemK produces a toxic effect in our wild type clinical strains.

To further confirm the relevance of PemK in pOXA-48 stability, we constructed a version of pOXA-48 lacking the toxin gene *pemK* (pOXA-48Δ*pemK*), and introduced it in *K. pneumoniae* KPN10, KPN11 and KPN18, and *E. coli* C288 and EC22. We performed plasmid stability essays for 5 days to determine the stability of both wt-pOXA-48 and pOXA-48Δ*pemK* in the different strain (Fig. 4A). Wild-type pOXA-48 was fully stable in *K. pneumoniae* and in *E. coli* EC22, but it was slightly unstable in *E. coli* C288. pOXA-48Δ*pemK* was surprisingly stable in *K. pneumoniae,* only starting to be lost in KPN11 and KPN18 after 4 and 3 days, respectively. Contrary, pOXA-48Δ*pemK* was very unstable in *E. coli,* being lost up to 90% in EC22. Both pOXA-48 and pOXA-48Δ*pemK* showed an erratic behaviour in *E. coli* C288, which might be attributed to the fact that the plasmid does not produce a high fitness cost in the strain, and therefore there is a small benefit associated with losing it (Fig. 1D). These results suggested that pOXA-48 segregation is more efficient in *K. pneumoniae* than in *E. coli*.

**Fig. 4.**
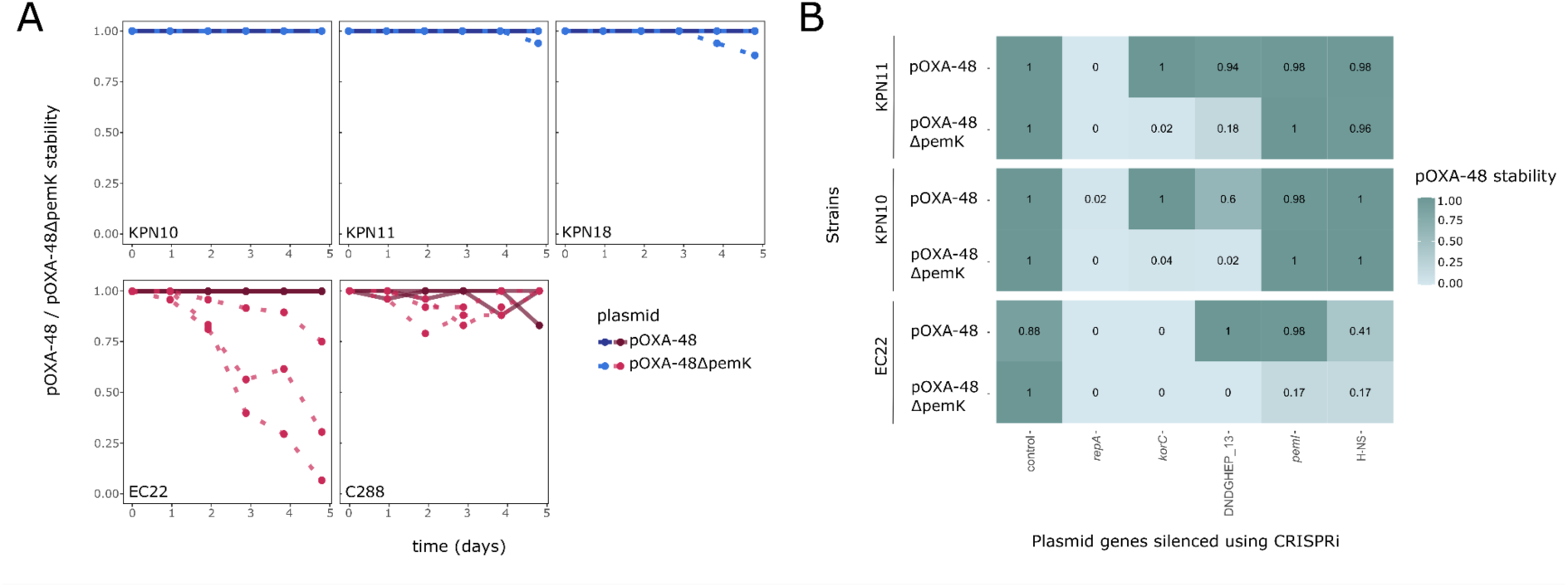
Role of PemK in pOXA-48 stability. The PemI-PemK TA system promotes pOXA-48 vertical transmission. (A) Stability of pOXA-48 (darker lines) and pOXA-48Δ*pemK* (lighter lines) in the different clinical strains over 5 days in LB culture (with a daily 1:100 dilution). (B) pOXA-48 and pOXA-48Δ*pemK* stability after 24 h of CRISPRi gene silencing of individual plasmid genes.

Finally, we decided to test the effect of silencing the three genes producing partial plasmid instability when silenced (*korC*, H-NS, and DNDJGHEP_0013-0014) in the pOXA-48Δ*pemK* plasmid, including the *repA* guide as control. Our hypothesis was that if PemK is eliminating the cells where pOXA-48 is lost, once the *pemK* gene is removed, we should be able to observe a decrease in plasmid stability associated with silencing these genes. We performed these experiments in *K. pneumoniae* KPN10, KPN11 and *E. coli* EC22. In line with our hypothesis, we observed an increase in plasmid loss associated with silencing these three genes (Fig. 4B). Again, the effect was stronger in *E. coli* than in *K. pneumoniae*. These results confirmed the relevance of *korC* and the DNDJGHEP_0013-14 operon in pOXA-48 stability, both in *E. coli* and *K. pneumoniae* clinical strains. The H-NS gene, on the other hand, seems to produce a species-specific effect, as suggested by our previous results (Fig. 3B).

Taken together, our results underline the key role of the PemI-PemK TA system promoting pOXA-48 vertical transmission in clinical enterobacteria.

### *bla*OXA-48 is responsible for pOXA-48 fitness costs

Only two pOXA-48 genes produce a significant fitness benefit when silenced in the absence of ertapenem, *repA* and *bla*_OXA-48_. The results for *repA* are puzzling, since we have demonstrated that *repA* silencing produces a rapid plasmid loss, which leads to a toxic effect. In fact, when looking at the CRISPRi screen results, *repA* is the only gene with significant effect that changes in sign over time. Silencing *repA* is associated with a fitness defect in the first time point and it becomes beneficial later on (SFig. 7). We comment on these results in more depth in the discussion section. On the other hand, previous reports have already indicated that *bla*_OXA-48_ expression is associated with a fitness cost in a clinical enterobacteria strain^26^. Therefore, we decided to study *bla*_OXA-48_ effects in more detail.

First, we correlated the CRISPRi score of *bla*_OXA-48_ with the relative growth of pOXA-48-carrying strains, obtaining a strong negative relationship (Fig. 5A, Pearson R-squared = −0.91, p-value = 0.0016, for *K. pneumoniae* and Pearson R-squared = −0.88, p-value = 0.02, for *E. coli*). This correlation was in fact stronger than the one obtained including the scores of all the genes in the CRISPRi screen (SFig. 3), as well as stronger than the correlation obtained with the subset of genes producing significant fitness effects (SFig. 8-9). Therefore, these results strongly suggest that the main contributor to pOXA-48 cost is the *bla*_OXA-48_ gene.

**Fig. 5.**
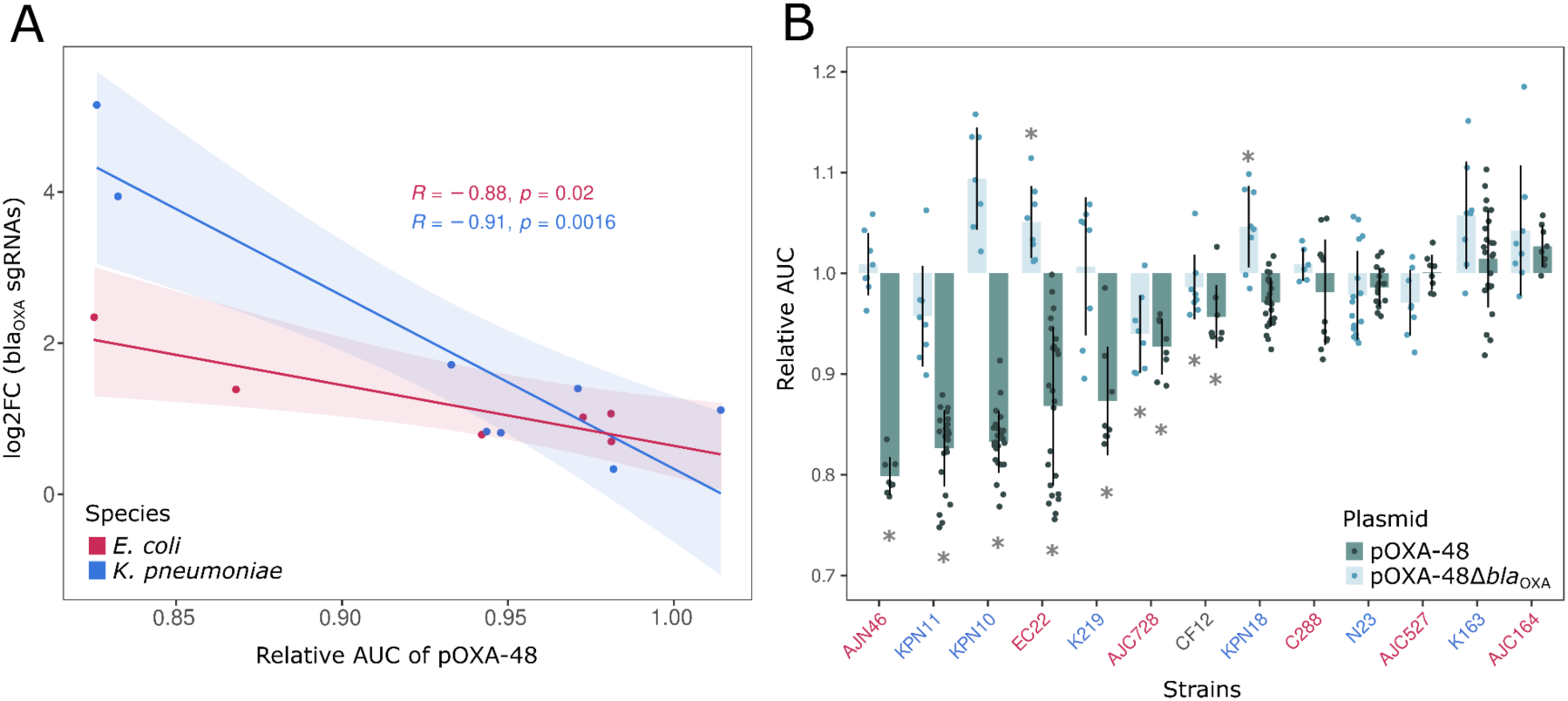
Role of *bla*_OXA-48_ in pOXA-48 fitness effects. The *bla*_OXA-48_ gene is responsible for pOXA-48-associated fitness costs. (A) Pearson correlation between the relative Area Under the growth Curve (AUC) of pOXA-48-carrying strains compared to their pOXA-48-free counterparts (shown in Fig. 1D) and the log fold change associated with silencing *bla*_OXA-48_ in the CRISPRi screen. Lines indicate the best fit for the correlation of each species. Shadowed areas indicate the 95% CI. Each color represents a species. (B) Plasmid cost associated with carrying pOXA-48 and pOXA-48Δ*bla*_OXA-48_ in different clinical strains of enterobacteria measured as relative AUC of pOXA-48-carrying compared to pOXA-48-free clones. Colours in legend are again indicative of species: in pink *E. coli,* in blue *K. pneumoniae,* and in black *C. freundii.* Asterisks indicate significant cost/benefit of either pOXA-48 or pOXA-48Δ*bla*_OXA-48_ in each of the strains (paired t-tests against each relative plasmid-free strain after Bonferroni correction; n = 8-16; p < 0.05).

To experimentally validate these results, we used a version of plasmid pOXA-48 with a small deletion affecting the *bla*_OXA-48_ gene (pOXA-48Δ*bla*_OXA-48_). This plasmid was recently recovered from a clinical strain and it does not produce a functional OXA-48 carbapenemase, but is otherwise isogenic to pOXA-48^26^. To minimise the potential overfitting caused by using the same strains throughout our study, we conjugated pOXA-48 and pOXA-48Δ*bla*_OXA-48_ in a second set of clinical enterobacteria including *Klebsiella* spp., *E. coli*, and *Citrobacter freundii* clones (S2 table). As previously described^12^, pOXA-48 produced variable fitness effects across the collection (average relative AUC= 0.938, SD= 0.075), causing a significant reduction in relative AUC in 7 out of 13 strains (paired t-tests against each relative plasmid-free strain after Bonferroni correction; n = 8-16; p < 0.05, Fig.5B). Interestingly, and in line with the hypothesis of *bla*_OXA-48_ being responsible for pOXA-48-associated fitness costs, pOXA-48Δ*bla*_OXA-48_ produced milder fitness effects in this collection (average relative AUC= 1.002, SD= 0.059), leading to significant costs in 2 of the strains and to significant fitness advantages in 2 other strains (paired t-tests against each relative plasmid-free strain after Bonferroni correction; n = 8-16; p < 0.05, Fig. 5B). These results confirm that the main cause of the fitness cost produced by pOXA-48 is the expression of the *bla*_OXA-48_ gene.

## Discussion

Unveiling the complex interactions between a plasmid and its bacterial hosts is essential for understanding plasmid ecology and evolution, and therefore for deciphering the dynamics of AMR in real-life scenarios, such as the hospital set-up or the patient’s gut. In order to better understand these interactions, in this work we study the fitness effect of each gene in pOXA-48, using 13 different clinically relevant clones of enterobacteria as hosts. To our knowledge, this is the first study to date to perform such an extensive screen in clinical strains, which allowed us to discover clinically-relevant interactions that could be missed by performing the experiments in laboratory strains. Our results showed that although pOXA-48 produces a wide range of fitness effects across the clinical clones (Fig. 1), the effects of silencing each plasmid gene are overall conserved (Fig. 2). This difference is driven by the fact that most of the gene-level fitness effects detected were “post-segregational killing” deleterious effects, associated with plasmid loss (Fig. 3-4). These effects emerge in the CRISPRi screen due to the silencing of genes associated with plasmid stability, but they do not necessarily contribute to the fitness effects of the wild-type plasmid with a correct partitioning. However, these results highlight the efficiency of plasmid-encoded mechanisms, such as the *pemI-pemK* TA system, promoting vertical plasmid transmission in clinical strains (Fig. 4). Finally, we could show that despite pOXA-48 carrying more than 90 genes and CDS, the fitness costs associated with this plasmid could be explained almost entirely by the expression of the carabapenemase gene *bla*_OXA-48_ (Fig. 5).

This study shows the potential of using CRISPRi screens to deepen our understanding of plasmid biology at a gene level. For example, we showed that apart from the previously known role in plasmid conjugation^36^, Pri plays a role in plasmid replication, likely as a primase (Fig. 3B-C). We also unveil the importance, particularly in *E. coli,* of three modulators of gene expression: KorC, an H-NS-like protein and DNDJGHEP_0014. Individually silencing the genes coding for those proteins led to plasmid loss in *E. coli* (Fig. 3B-C). The importance of KorC and DNDJGHEP_0014 (*orf20*) for pOXA-48 stability was also observed in *E. coli* MG1655 in an elegant parallel study by Baffert *et al* working with a transposon insertion library of pOXA-48^37^. The specific role of these three genes will be explored in more detail in the future.

We also show that, as previously described^38^, the PemI-PemK TA system promotes the vertical transmission of pOXA-48 to cell progeny. However, we observed that PemI-PemK role in vertical plasmid transmission varied among clones. pOXA-48Δ*pemK* was stably transmitted to the progeny in the 3 *K. pneumoniae* strains tested but not in 2 *E. coli* strains (Fig. 4A). Given that our data indicated that PemK is toxic both in *K. pneumoniae* and *E. coli*, these results suggest that plasmid segregation may be more efficient in the first species, which could help to explain why pOXA-48 is more strongly associated with *K. pneumoniae* than *E. coli* in the clinic^30^. We hypothesise that this higher stability could be explained by the fact that pOXA-48’s plasmid copy number is higher in *K. pneumoniae* than in *E. coli* (*K. pneumoniae,* mean = 2.21 copies; *E. coli* mean = 1.41), as we recently showed^7^. Alternatively, this difference could be due to the partition machinery being more efficient in the first species than the latter, which would suggest that pOXA-48 is better adapted to the *K. pneumoniae* than *E. coli*. Finally, it could also be possible that additional TA systems in pOXA-48 may explain the differences in plasmid stability between species. In fact, Baffert *et al.* just reported the existence of a second TA system in pOXA-48, composed of a toxin, DqlB, and a non-coding RNA *trans*-acting antitoxin, *agrB*^37^. This system had not been previously annotated in pOXA-48 due to its small size, and is located in the intergenic region between *repA* and *trbB*. Because of the lack of annotation, we did not design guides specifically targeting this region in the OXAlib. However, three of our *repA-trbB* intergenic guides target the antitoxin *agrB* (SFig. 10A). In line with the results from Baffert *et al.*, we observed a dramatic decrease in the log2FC of the guides silencing the antitoxin throughout our CRISPRi screens (SFig. 10B-C), supporting the role of TrbB-*agrB* as a second TA system in pOXA-48. The effect associated with silencing *agrB* was however similar for *K. pneumoniae* and *E. coli* strains, so it did not help to explain the higher stability of pOXA-48 in *K. pneumoniae*.

In our CRISPRi screens only two genes produced a fitness benefit when silenced in the absence of antibiotic selection: *repA* and *bla*_OXA-48_. The results for *repA* are difficult to explain, and highlight some of the limitations of our set-up. On the one hand, it is possible that a small subpopulation of the cells that carried a *repA* guide survived an early pOXA-48 loss in the experiment, overcoming the PemK and TrbB toxic effect. This subpopulation would be free of the cost associated with pOXA-48 carriage and could therefore proliferate in the pool of cells. This would explain the rapid initial decline of guides targeting *repA* in the population, and their subsequent enrichment in later time-points (SFig. 7). On the other hand, it has been shown that weak and unpredictable CRISPRi gene repression can arise from targeting CDSs under negative regulatory feedback loops^39,40^. Rep proteins are known to follow this type of self-regulation and this could also contribute to the seemingly erratic behaviour of guides targeting the *repA locus*^41^.

The main findings of this work is the fact that the gene encoding for the β-lactamase OXA-48 is the main contributor to pOXA-48’s fitness cost across clinical strains (Fig. 5). β-lactamases had already been shown to have an effect on bacterial cell physiology and lead to a decrease in fitness^25^. These effects may have different origins. First, they could be produced by the β-lactamase’s signal peptide^42^, responsible for folding, stability and translocation of the protein to the periplasm^43^. By leading to a host-specific inefficient processing of the protein and subsequent envelope stress, the signal peptide effectively dictates the host-range of some β-lactamases^44^, which could explain why plasmids encoding for these genes are more costly in some hosts than others. Second, the fitness costs associated with β-lactamases have also been associated with their residual DD-endopeptidase activity, which leads to the hydrolysis of the peptide-cross bridge and alters the structure and stability of the cell wall^45^. This means that carrying plasmids such as pOXA-48 can lead to a severe destabilisation of the bacterial envelope which can have downstream repercussions, such as fitness costs and even collateral sensitivity to other antibiotics^46,47^. This also paints a somewhat hopeful picture about the dissemination of these plasmids in the population: since the main source of cost of the plasmid is the AMR gene itself, in the absence of antibiotic treatment the cells will be under pressure to either lose the plasmid or mutate *bla*_OXA-48_. The latter was already observed in a previous study analysing within-patient evolution of pOXA-48, where a small deletion affecting the *bla*_OXA-48_ gene and abolishing the AMR phenotype was associated with reduction of plasmid cost in the gut microbiota of a hospitalised patient^26^. Lastly, this also opens a gateway to explore new ways to exploit these changes in the cell envelope as potential new antimicrobial strategies, ones highly specific to cells carrying genes within the β-lactamase family^46,48^.

In summary, our study highlights the role of different plasmid-encoded genes in the biology of pOXA-48 and opens new doors to explore alternative ways to fight AMR in enterobacteria. Our results also highlight the usefulness of using high-throughput techniques to explore previously unknown biological phenomena at a genetic level. And lastly, given that previous works indicate that both the fitness and transcriptomics response to carrying plasmids are highly host-specific^21,49,50^, our study paves the way to performing genome-wide CRISPRi screens in order to dissect plasmid-bacteria interactions in clinical bacteria. Set-ups such as this, although complex and labour-intensive, will help us unveil aspects of plasmid biology with applications in real-life scenarios.

## Methods

### Bacterial strains and culture conditions

All clinical strains used in this study were obtained from the previously characterised R-GNOSIS collection of over 28,000 rectal samples collected from patients admitted to the Hospital Universitario Ramón y Cajal in Madrid between the years of 2014 and 2016 (R-GNOSIS-FP7-HEALTH-F3-2011-282512)).

For the CRISPRi screens, a subset of these strains was selected based on previous works. First, a collection of strains that did not carry pOXA-48 naturally and where pOXA-48 transconjugants had been obtained^32^, and, secondly, a collection of strains that carried pOXA-48 naturally and where pOXA-48 had been successfully cured^26^. Out of these two collections, strains were selected based on sensibility to apramycin - antibiotic resistance gene used for selection of the OXAlib library -, conjugation rate efficiency, belonging to a sequence type of clinical relevance, their plasmid profile and, lastly, activity of dCas9 - tested by measuring cell death by silencing the essential gene rpsL, as in Cui *et al*.^33^ (Supplementary Methods, SFig. A)). The resulting selection comprises 14 strains, 8 *K. pneumoniae* strains and 6 *E. coli* strains (including the lab strain J53) (S2 table).

The variant of pOXA-48 used in this work is pOXA-48_K8 (accession number MT441554), and its sequence and complete genomes of the strains used in this study were obtained by the lab in previous works, available under the BioProjects PRJNA641166 and PRJNA803387 in the SRA repository of the National Center for Biotechnology Information (NCBI).

All experiments were performed in Lennox lysogeny broth (LB), supplemented with 15 g/L of agar when indicated. The different antibiotics used throughout the study were amoxicillin / clavulanic acid at a proportion of 5 mg amoxicillin / 1 mg clavulanic acid, ertapenem, and apramycin. 50 µM of 2,4-diacetylphloroglucinol (DAPG) was used for induction of dCas9.

### Calculation of pOXA-48 fitness effects in different clinical strains

The fitness effect associated with pOXA-48 in each strain was calculated by comparing differences in the Area Under the Curve (AUC) of isogenic plasmid-carrying and plasmid-free clones, as described in DelaFuente *et al*^51^. Raw data can be found in the supplementary materials (S4 table).

### Designing the OXAlib library

dCas9 is reprogrammed to target a specific position using guide RNAs (sgRNAs), the design of which needs to follow a specific set of rules in bacteria^52^. (i) First, it has been shown that the specific sequence of the sgRNA has an effect on the activity of dCas9. (ii) Due to design limitations intrinsic to the CRISPRi technique, sgRNAs targeting coding regions have to be limited to the coding strand. The rules for efficiently targeting non-coding regions of DNA are not known, and therefore sgRNAs targeting those regions should be designed to target both strands of DNA. (iii) If dCas9 is directed to a gene within an operon, this binding will have a polar effect on the downstream genes. (iv) It is also important to avoid potential off-targets resulting from a permissiveness of dCas9 to bind to similar regions of DNA to *bona fide* targets.

The OXAlib library was designed using the CRISPR@Pasteur design tool which is trained on *E. coli* activity data, and which has been successfully used to design such libraries in different species of Enterobacteriaceae^52^. The tool was adapted to our setup (looking for targets on a plasmid and for off-targets on a set of genomes - see Code Availability). Using this tool, the 5 best-performing sgRNAs were selected for each gene in pOXA-48_K8 (called pOXA-48 in this paper). sgRNAs targeting both DNA strands were also designed against intergenic regions of the plasmid (up to 5 targeting each strand, depending on the size of the intergenic region). The logic behind targeting both strands in intergenic regions is to attempt to capture effects of targeting any possible genetic elements present in the regions. 20 non-targeting control sgRNAs were also designed, amounting to a total of 568 sgRNAs (S1 table).

Guides were selected with the previously described sorting strategy and ordered for synthesis as ssDNA oligos from Twist Bioscience. The sequence of each individual oligo is as follows: 5’-ACAGCTAGCTCAGTCCTAGGTATAATACTAGT-(20nt guide sequence)-GTTTTAGAGCTAGAAATAGCAAGTTAAAATAAGGCTAGT-3’.

### Cloning the OXAlib library

The ssDNA oligo pool was amplified using primers AC74 and AC75 (S2 Table) for subsequent cloning into pFR56apm^21^. pFR56apm is based on pFR56^53^, with an apramycin resistance gene as a selection marker. This antibiotic permits selection of the vector in all the different clinical strains (which due to their plasmidsome are naturally resistant to several antibiotics). Amplification was performed using New England Biolabs’ Phusion High-Fidelity DNA polymerase, with the HF buffer, DMSO and the recommended protocol. To minimise PCR amplification biases of specific oligos in the pool, the reaction was limited to 9 cycles (1 cycle for generating dsDNA oligos and 8 cycles of amplification), and performed with 1ng of template and 50 pmol of each primer. The resulting product was gel purified. The pFR56apm vector was digested with *BsaI* and gel purified. The OXAlib plasmid library was assembled using the Gibson Assembly method^54^, using New England Biolabs’ NEBuilder HiFi DNA Assembly, 200 ng of total DNA and a ratio of 1:5 vector:insert. Cloning was performed in *E. coli* MG1655. Quality control of the final OXAlib library can be found in the Supplementary Methods.

### Conjugating the OXAlib library into clinical strains

pFR56apm is a mobilisable plasmid - meaning it can be conjugated to compatible hosts, but requires expression of the conjugation machinery in *trans*. The OXAlib library was minipreped from *E. coli* MG1655 and transformed (with a coverage >1000x of OXAlib) into *E. coli* ß3914ø + pTA-MOB^55,56^. This strain is auxotrophic for diaminopimelic acid (DAP), allowing for its counter-selection as a donor strain.

Conjugation of the OXAlib from ß3914ø to the different strains was performed by mixing 10^8^ donor cells with 10^8^ of recipient cells (ensuring a coverage >1000x of OXAlib) for 3 h. After conjugation, the cells were selected in apramycin (50µg/mL) and let grow overnight at room temperature. Absence of DAP counter-selects against ß3914ø. Conjugation efficiency was assessed using serial dilutions and a good coverage of the OXAlib library achieved for each strain (S1 table).

### CRISPRi screens

#### Experiment

First of all, it is crucial when working with CRISPRi screens to ensure that each step guarantees a good coverage of the library (*i.e.* that no individual guides are lost from the population due to over-dilution). To ensure this, we always work with an excess of cells of at least 1000x the size of OXAlib.

Strains carrying the OXAlib library were diluted 1:1000 in fresh LB / LB supplemented with ertapenem (subinhibitory concentrations of the antibiotics were used, Supplementary Methods, S1 table). After 1 h, DAPG (50 µM) was added to the cultures for dCas9 induction. Every 8 h, a 1:1000 dilution was performed in fresh LB + DAPG / LB + DAPG + ertapenem. Prior to each dilution, OD was measured (to calculate number of generations). At 24, 48 and 72 h, apart from the dilution and OD measurement, a miniprep was performed using 10 mL of culture. The resulting pool of plasmids correspond to the timepoints at ∼25, ∼55 and ∼85 generations. All screens were performed in triplicate (3 biological replicates), in aerobic conditions, at 37°C, with shaking (250 r.p.m.).

#### Illumina sample preparation and sequencing strategy

Library sequencing was performed as previously described^33^, with some small adaptations. The PCR primers used for the nested PCR can be found in the supplementary materials (S2 table). To maintain good coverage of the OXAlib library, 100ng of plasmid miniprep were used as template for each reaction. To minimise PCR biases, 19 cycles were used in the 1st PCR, and 13 cycles on the 2nd. Each replicate was given a different i5-i7 index combination, achieved by amplification with specific pairs of primers. Lastly, Illumina sequencing quality depends strongly on the variability of the regions sequenced. To bypass variability issues arising from amplicons sharing a large 5’ identical sequence, 3 different staggered primers were used in the 1st PCR.

Each sample was quantified using a bioanalyzer. 150 ng of each sample were pooled together and sequenced using the standard protocol of a NextSeq 500 benchtop sequencer (Illumina).

### Data analysis

#### Quality control of Illumina sequencing

First, a QC of the raw Illumina data sequences (55 bp amplicon sequencing) was performed using FastQC v.0.11.9^57^ and MultiQC v.1.11^58^. After confirming high quality per base score in the guide region (mean Phred score > 30), the sequences were used in downstream analyses.

#### Guide count per sample

For getting each individual guide count in each sample, each fastq file was parsed using Bash (see Code Availability). Briefly, knowing the sequence of the promoter preceding the guides in each read, 15 bp of it were used to match the reads position in which the guide began, and the following 20 bp (i.e., the sgRNA) were extracted. For each sample, the number of times each individual guide appeared was calculated using Bash commands. The list of guides was sorted, each unique occurrence was counted, and the output was reformatted to save each strain guides count in an individual csv file. Within the individual csv counts files, the guides with less counts than 5 for timepoints 0 and with less counts than 2 for the rest of the timepoints were filtered. The results of all the samples were merged in the same table, and only those guides present in the plasmid library were kept. The sequencing coverage for each guide was calculated as the median of the counts of guides (1 sgRNA per read; avoiding outliers from guides counted in excess) multiplied by 5 (as there are 5 guides per gene). The counts for the three replicates per time point were merged after checking the reproducibility of the replicates in each condition (LB+DAPG or LB+DAPG+ertapenem).

#### Gene score calculation

For calculating the gene scores (median log2 fold change by gene) a script which took the table with the replicates merged by sample, and calculated the log2FC for each gene, then centering the values by removing the median of control guides, was written based on the code of Rousset *et al*^53^. (https://gitlab.pasteur.fr/dbikard/ecocg/-/blob/master/Notebook.ipynb).

#### Phylogenetic analysis

The phylogenetic distances between the different strains were calculated using Mashtree v.1.2.0^59^ (NJ method) and visualised using the iTOL web server (https://itol.embl.de/). Phylogenetic information was used to represent the strains clustered by relatedness in the heatmap (Fig. 2A).

#### Conservation of CRISPRi screening results in different strains

To analyse whether the effect of the OXAlib library on the strains tested was conserved, we studied the similarity between strains gene scores at the end of the screening. We calculated the correlation values (Pearson correlation coefficients) between the strains based on all the gene scores at the end of the screening for each condition by using the corrplot package v.0.95 in R. Additionally, we compared the genes whose silencing had the most notable effect in each strain. To do so, we listed the significant genes and intergenic regions for each strain and condition calculated from individual permutation tests for either LB+DAPG or LB+DAPG+ertapenem (see following Methods section). With these, we measured the Sørensen-Dice Similarity Coefficient (SDC) for all pairwise comparisons between strains of the same species screened in the same condition. The SDC was calculated using the following formula:

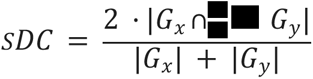

Where |*G_x_*| and |*G_y_*| are respectively the number of significant genes in strain X and Y (i.e. the number of elements compared from each strain), and |*G_x_* ∩ *G_y_*| is the number of common significant genes between X and Y strains in the given condition (i.e. full set of significant genes for both strains in either ertapenem or no antibiotics).

#### Permutation tests for gene importance

For checking statistically significant differences from the score of each gene from the distribution of gene scores in the CRISPRi screen, a permutation test for all the strains considered together per each condition (LB+DAPG or LB+DAPG+ERTA) was carried out. Briefly, the mean of each gene score was calculated considering all the strains from the screens (n = 14). The difference of this value and the mean of the scores from the rest of the group was assessed as the statistics of interest in the permutation test. The same statistic was calculated for 100000 iterations (i.e., minimum computable p-value = 0.00001) taking random samples (n = 14) in each iteration. This number of permutations was selected to get more precise probabilities, as the p-values of some genes resulted marginally significant when performing the permutation tests with fewer repeats (1000 - 10000). Lastly, the number of times that the permuted statistic was more extreme than the mean of each gene (i.e. the p-value) was calculated. The same test was conducted for each species, as well as for each strain, for both experimental conditions. Finally, the p-values were adjusted by FDR. To report the statistical results from the permutation tests as well as the CRISPRi screening results, Volcano Plots were represented to simultaneously show each gene p-value and its score using the R package EnhancedVolcano v.1.18.0. As in this type of screen, most genes showed scores close to 0. Thus, a log2FC low threshold (i.e. +-0.5) was set, close to the centre of the distribution. For the p-value threshold, the default one (10e-06; i.e. p-value < 0.05) was selected.

#### Ridge regression model for plasmid cost

To simultaneously consider the contribution of multiple genes to pOXA-48 cost, a multiple regression model with Ridge regularisation was developed using the glmnet v.4.1-7 R package separately for both *E. coli* and *K. pneumoniae*. First, feature selection of all the genes and intergenic regions tested was performed, as there were 8 observations of plasmid cost for *K. pneumoniae* and 6 observations of plasmid cost for *E. coli* (n=strains), and 106 explanatory variables (n=genes & intergenic regions included in CRISPRi screen). Then, only those genes with significant p-values in the permutation test performed for each of the species (i.e. expected to be more informative) in absence of antibiotics during the CRISPRi screen were included in the model. From these, intergenic regions were kept out, as their effect introduced multicollinearity issues by showing similar behaviour as their surrounding genes in the CRISPRi screening (e.g. *bla*_OXA-48_ intergenic surrounding regions). Multicollinearity between explanatory variables was assessed after feature selection using the R package corrplot v.0.95. Due to the limited number of observations for each species, the models were trained using Leave-One-Out Cross Validation. Moreover, to include all the genes resulting from the feature selection process, Ridge regularisation was selected, as the regression coefficients are not shrunk to 0. Optimal parameters from the trained models were selected to then build the final models. The ridge coefficients for each gene, and the R-squared for each species were calculated from the final models tuned with the optimal parameters (lambda [λ]; SFig. 8).

#### Plasmid cost based on guide distribution

To infer the plasmid cost in each strain, a proxy based on the general effect of blocking the plasmid genes was employed. The effect of each plasmid gene on bacterial fitness is expected to be reflected on the CRISPRi screen in absence of antibiotics, as guides which block costly genes that cause deleterious effects are expected to show higher scores, and vice versa. Then, if the plasmid is highly costly in a strain, the general effect of blocking its genes should reflect this. Thus, the general effects of blocking the plasmid genes were correlated with the relative fitness of plasmid-carrying strains. The difference between the mean effect of blocking pOXA-48 genes (mean CRISPRi gene scores for each strain) and the mean effect of the control guides, which are supposed to have no effect, was calculated. Then, these values were correlated (Spearman method) with the fitness effect of pOXA-48 in each strain.

### Validation of CRISPRi screen’s results

#### Strain selection

From the initial set of strains used in this study, two *E. coli* (C288, EC22) and three *K. pneumoniae strains* (KPN10, KPN11 and KPN18) were chosen for validation of results based on the overall strongest fold-changes observed.

#### Cloning of individual sgRNAs in pFR56apm and conjugation into clinical strains

Cloning of individual sgRNAs (S1 table) in pFR56apm was performed using Golden Gate assembly as in Rousset *et al*.^53^ All cloning was performed in *E. coli* MG1655 and followed by conjugation into the clinical strains as described previously.

#### bla_OXA-48_ associated fitness effects

To validate the results associated with silencing the gene *bla*_OXA-48_, growth curves were performed to study plasmid fitness effects as explained before in this method section. This time, a second set of strains was selected from the RGNOSIS collection, to avoid overfitting to the original strains used in the screen. These strains were once again selected based on the fitness effects associated with the plasmid, as described in Fernández-Calvet *et al*.^12^ A variant of pOXA-48 lacking a complete *bla*_OXA-48_ was identified previously in the lab^26^. This variant was conjugated on the cured version of the strain, obtaining three isogenic set-ups for each strain: the wild-type carrying pOXA-48, a pOXA-48-free cured version, and a pOXA-48Δ*bla*_OXA-48_ transconjugant.

#### Plasmid stability

Plasmid stability of pOXA-48 and pOXA-48 versions was measured by growing cells carrying the plasmid in LB over subsequent periods of 24 h. After each cycle, cells were plated in LB, guaranteeing the isolation of individual colonies. Cells were then streaked in presence and absence of ertapenem using replica plating. Percentages of plasmid loss / plasmid presence were calculated by comparing growth in presence and absence of selection. Each experiment was performed in triplicate. Number of 24 h cycles used in each experiment are indicated throughout the text. To statistically analyse whether the stability of pOXA-48 was significantly affected in at least one strain when silencing individual genes (Fig. 3B), we performed a one-tailed z-test for each guide. Briefly, we calculated the maximum deviation from the control group (guide “control”) per individual guide, from which we inferred their Z-score. Finally, we corrected the p values by Bonferroni adjustment (n = 8).

#### qPCR for PCN calculation

Changes in pOXA-48’s PCN were calculated for the different clinical strains carrying pFR56apm programmed to silence different plasmid genes. In order to do so, the different strains were grown in triplicates in LB supplemented with apramycin 50 µg/mL and DAPG 50 µM for 24 h. 50 µL samples were taken of each replicate, centrifuged to collect the pellet and boiled for 10 min. 2 µL of each boiling reaction were used per qPCR reaction, using the NZYSupreme qPCR Green Master Mix (2X), ROX plus kit from NZYTech. Targeted plasmid and chromosome genes were *bla*_OXA-48_ (amplicon size 100bp; efficiency 107,47%) and *dnaE* (chromosomal gene with 1 copy, amplicon size 200bp, efficiency 105.79%), respectively. Efficiency was calculated (taking into account amplicon size difference) as in San Millán *et al.*^60^ The amplification conditions were: 5 min of initial denaturation (95°C), followed by 30 cycles of 15 s denaturation, 30 s annealing (55°C) and 30 s extension (60°C). The relative PCN was calculated as in San Millán *et al.*^60^ Raw data for qPCR can be found in the supplementary data (S4 table).

#### Lambda-Red recombination for construction of deletion mutants

A version of pOXA-48 lacking *pemK* was built using lambda-red recombination^61^. The recombination cassette with extremes matching the beginning and end of *pemK,* and a kanamycin gene flanked by FRT sides was amplified from pKD4, as described in Datsenko *et al*.^61^ Primers used for construction of the cassette and validation of the construct can be found in the supplementary data (S2 table).

## Data availability

The sequences generated during this project are available under the BioProject ID PRJNA1203587 in the SRA repository of the National Center for Biotechnology Information (NCBI).

## Code availability

All the code developed for the analyses included in this work can be found at the Github repository https://github.com/jorgEVOplasmids/CRISPRi_pOXA48.

## Acknowledgements

We thank the R-GNOSIS group, the volunteers and the medical staff from the Hospital Universitario Ramón y Cajal (Madrid, Spain) involved in the sample isolation process. We thank David Bikard and the Institut Pasteur for plasmid pFR56. We thank the *Unidad de Genómica del Parque Científico de Madrid* for helping with the Illumina sequencing. This work was supported by the European Research Council (ERC) under the European Union’s Horizon Europe research and innovation programme (ERC-2022-CoG Project 101086992 – PLAS-FIGHTER) and by MCIN/AEI/10.13039/501100011033 and the European Union NextGenerationEU/PRTR (Project PCI2021-122062-2A). A.CV. was funded by an EMBO postdoctoral fellowship (ALTF 322-2022). We acknowledge financial support from the Spanish State Research Agency, AEI/10.13039/501100011033, through the “Severo Ochoa” Programme for Centres of Excellence in R&D [SEV-2013-0347, SEV-2017-0712, CEX2023-001386-S].

## Author contributions

A.CV., J.SD. and A.S.M. conceptualised the study. A.CV., J.SD., A.BM. and A.S.M. designed the methodology. J.SD. performed the bioinformatic analyses. A.CV. A-BM. and C.C.R. performed the experiments. A.CV., J.SD. and A.S.M. contributed to data analysis. A.S.M. was responsible for funding acquisition and supervision. A.CV., J.SD. and A.S.M. wrote the original draft and undertook the reviewing and editing process. All authors supervised and approved the final version of the manuscript.

## Supplementary Material

### Supplementary Methods

#### Quality control of the OXAlib library

Quality of the OXAlib library was assessed by MinION sequencing. Analysis of the library pool revealed the OXAlib to be composed of 1868 individual guide sequences, with 564 out of the 568 designed guides present. These differences between the designed and the obtained library are explained by errors during the synthesis of the initial oligo pool. The distribution of these guides in the population revealed that 87.5% of the pool corresponds to the correct 564 guide sequences. Therefore, the quality of the OXAlib library was deemed sufficient for its use in this study.

#### MIC determination and antibiotic concentration for CRISPRi screens

The Minimal Inhibitory Concentration (MIC) of ertapenem in liquid was calculated for each isogenic pOXA-48-carrying and plasmid-free couple of strains. 1:1000 dilutions of overnight cultures of each strain were diluted in LB supplemented with doubling concentrations of ertapenem (starting at 0.125 up to 4 µg/mL) and grown overnight. The MIC for ertapenem corresponds to the lowest concentration of antibiotic that does not allow growth after the overnight incubation (S1 table).

In order to select the sub-inhibitory concentrations or ertapenem for each strain to use during the CRISPRi screens, we decided to work with the highest sub-inhibitory concentration of antibiotic in liquid that would not lead to a growth defect in the plasmid-free strain. This creates a set-up where the cell does not suffer strong metabolic changes associated with the stress of an antibiotic whilst still generating a small benefit of carrying the plasmid (S1 table).

#### dCas9 activity in strains of work

The activity of dCas9 expressed from pFR56apm was tested in all strains of work by using the CRISPRi machinery to block the essential gene *rpsL.* Individual sgRNA against *rpsL* were cloned using the Golden Gate method (see main Methods section). Because a single sgRNA against all strains of work was not possible, we designed three individual sgRNAs against *rpsL,* each working in different sets of strains: rpsL (against all strains of *K. pneumoniae),* rpsL3 (against *E. coli* EC03, EC22, EC23, C288 and J53) and rpsL4 (against *E. coli* EC05, EC08 and J53) (Table S1)

#### pFR56apm mobilisation during screens

One big concern during the CRISPRi screens was that the different sgRNA-carrying pFR56apm could be mobilised between cells during the screening protocol. This would be problematic as it would add a strong noise to the dataset.

This would be very unlikely. First, because the experiment is performed in shaking conditions, which make conjugation very difficult. And second, as explained in the main Methods of this paper, because pFR56apm can’t be mobilised to a new cell in the absence of a second plasmid that provides the necessary conjugation machinery. An analysis of the plasmidsome of the different clinical strains used in this study showed that, theoretically, none of them carried conjugation genes able to mobilise pFR56apm. However, we wanted to make sure whether conjugation of pFR56apm did not take place.

To do this, we tested the different clinical strains’ sensitivity to sodium azide, showing all of them except *E. coli* J53 to be sensitive to sodium azide 100 µg/mL.

Then, we mimicked the conditions of the CRISPRi screen experiment, individually mixing 1:1000 dilutions of overnight cultures of J53 not carrying pFR56apm, with 1:1000 dilutions of overnight cultures of the rest of the clinical strains carrying pFR56apm with a non-targetting sgRNA. We grew the cultures in LB, at 37°C with shaking (250 r.p.m.). After 6 h of growth, we plated 50 µL of the culture in LB supplemented with sodium azide (100 µg/mL) and apramycin (50 µg/mL). No growth was observed in any of the plates, showing that conjugation of pFR56apm does not take place in the conditions of the CRISPRi screen.

#### Stability of pFR56apm and pOXA-48 during screens

Plasmid stability of pFR56apm and pOXA-48 were measured during the screens following the same protocol as described in the main methods of this paper, with the following alterations: a single time-point was assessed (72 h, the end of the screen experiment).

### Supplementary Figures

**SFig. 1.**
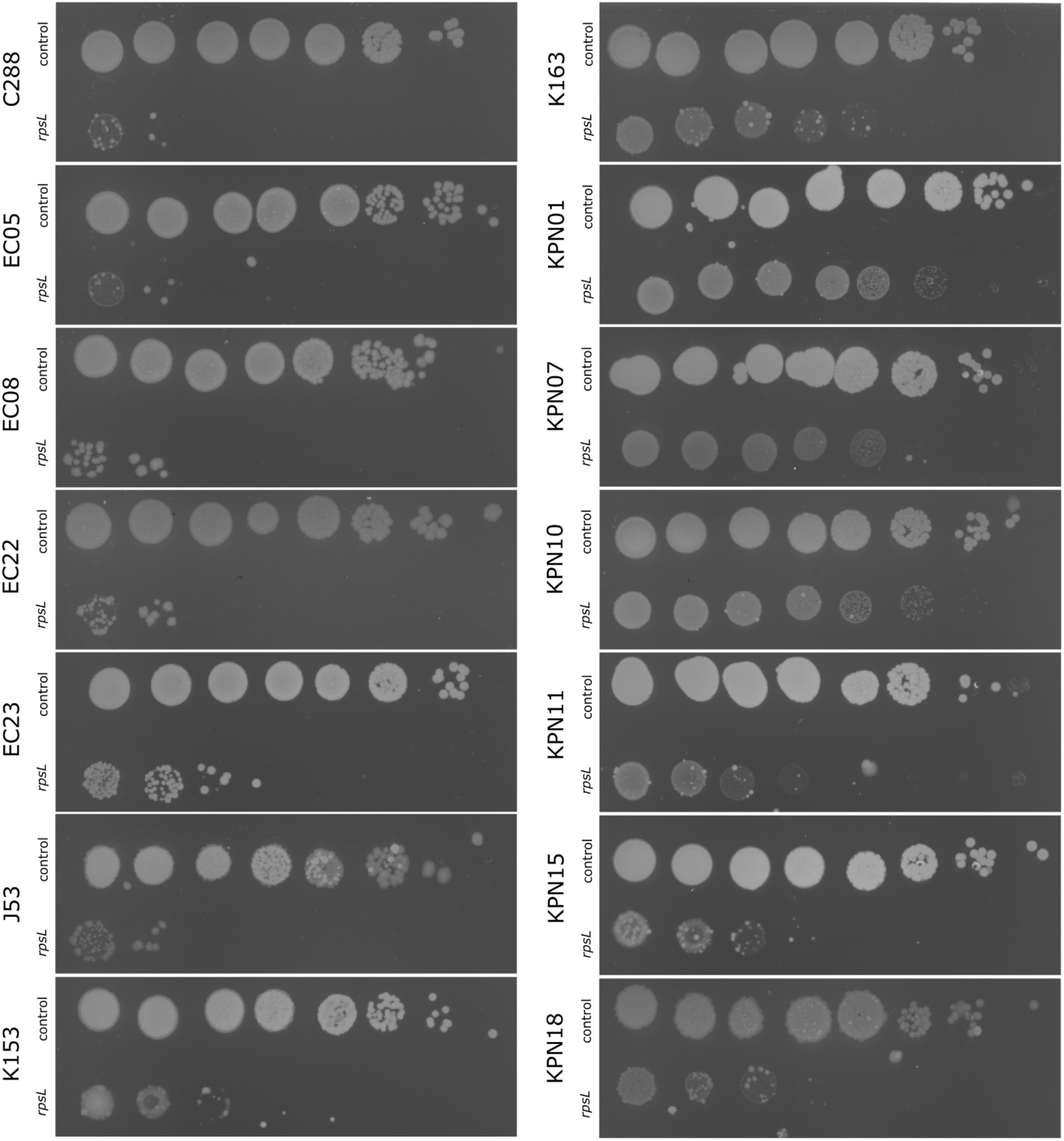
Efficiency of CRISPRi silencing machinery in the different clinical strains. Growth on LB agar after an overnight culture of serial dilutions (1/10) of different strains carrying the CRISPRi machinery programmed with a non-targeting control (should not lead to killing) or with an sgRNA against the essential gene *rpsL* (should lead to killing).

**SFig. 2.**
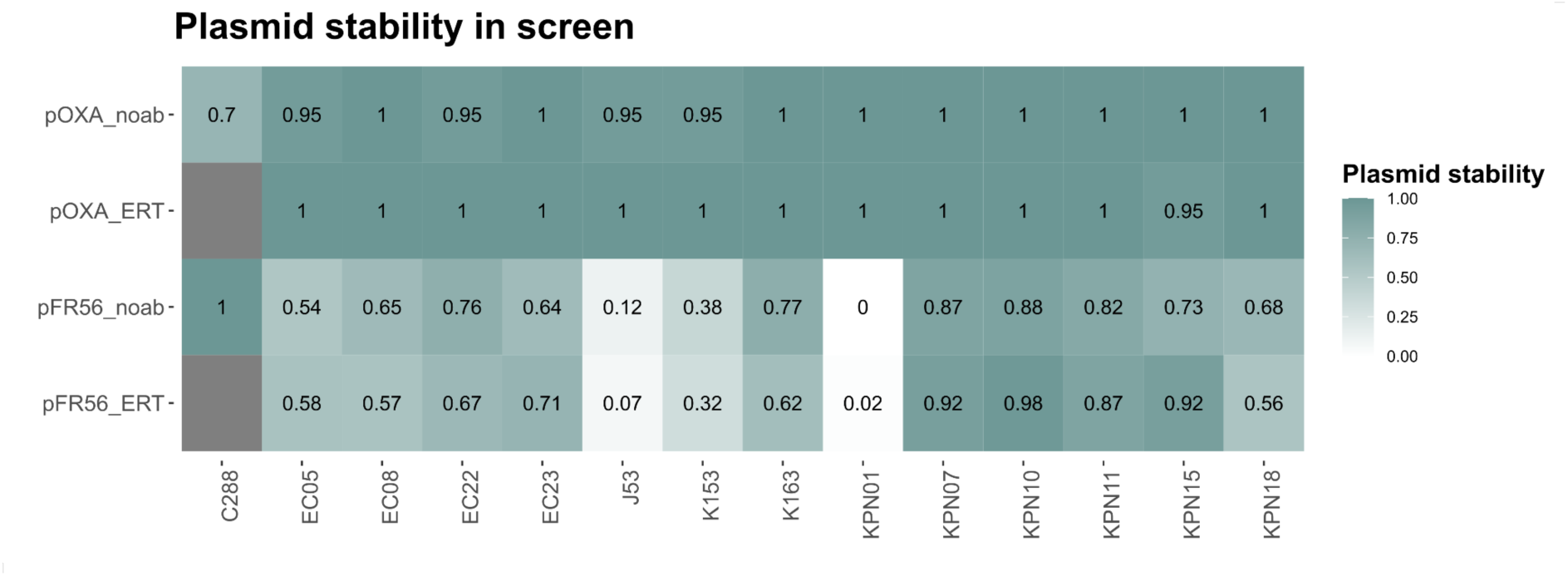
Stability of pFR56apm and pOXA-48 after 72 h of CRISPRi screens in the different clinical strains. Stability of both pOXA-48 and pFR56apm were assessed after the CRISPRi screens performed in presence of subinhibitory concentrations of ertapenem (ERT) or no antibiotic (noab). We were not able to obtain data for strain C288 in the presence of Ertapenem.

**SFig. 3.**
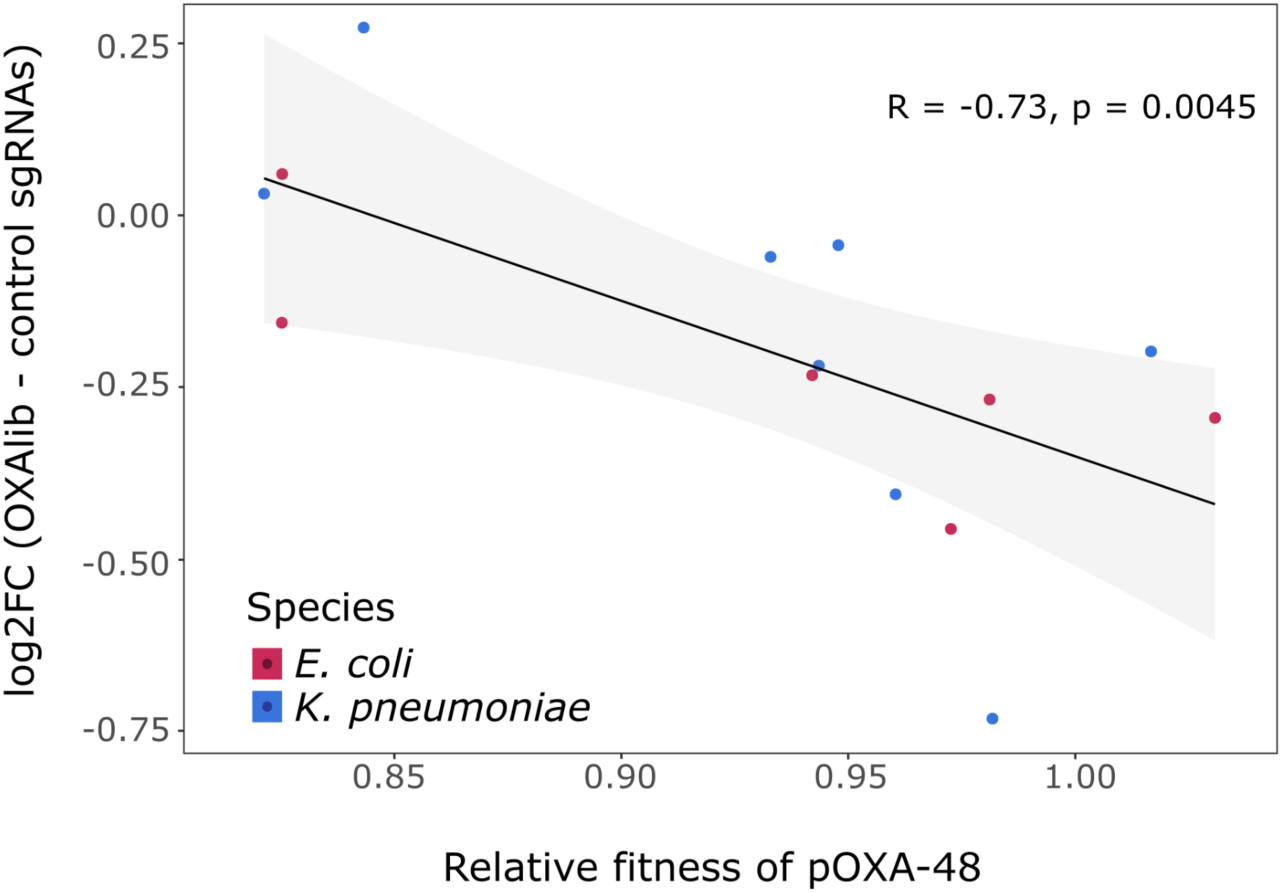
Spearman correlation between pOXA-48 fitness effects in the different strains under study (shown in Fig. 1D) and the fitness effects of all individual plasmid genes measured in the CRISPRi screen. pOXA-48 fitness effects were calculated as the Area Under the Growth Curve (AUC) for each bacterial strain carrying the plasmid (n = 16) relative to the mean AUC of the isogenic plasmid-free strain. The effects of all individual plasmid genes were calculated as the difference between the mean effect of blocking pOXA-48 genes (mean CRISPRi gene scores for each strain) and the mean effect of the control guides (see Methods “Plasmid cost based on guide distribution” section) for each strain. Black line shows the best fit for the Spearman correlation between both variables, whereas the grey shadow indicates the 95% confidence interval. Point colors represent the species of each bacterial strain.

**SFig. 4.**
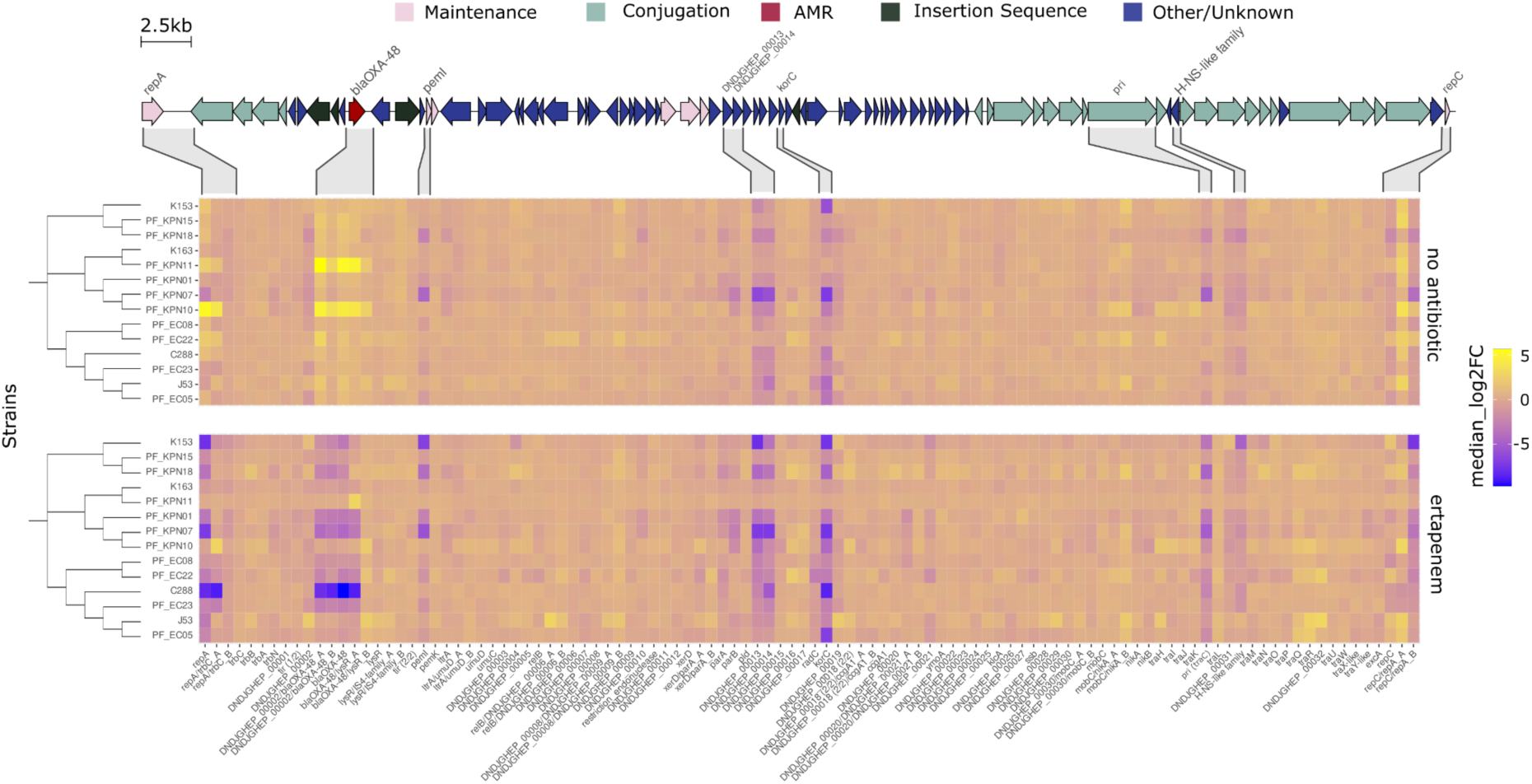
CRISPRi screens results for each pOXA-48 gene - fully annotated. Heatmap of CRISPRi gene scores at the end of the screening (t = 72 h, ∼85 generations). Fitness effects associated with silencing each individual gene / intergenic region in the absence of selection (top panel) and in the presence of ertapenem (bottom panel). The score for each gene / intergenic region corresponds to the median log_2_ fold-change of the sgRNAs targeting the element in the population (see Methods). Blue shades indicate those genes which silencing is detrimental (i.e. they are beneficial in that condition), whereas yellow shades correspond to genes which silencing is beneficial during the experiment (i.e. they are costly in that condition). Coding regions of pOXA-48 are shown with different colours indicating common functional clusters as shown on top. Each gene / intergenic region is indicated in the bottom of the figure, and those which showed a significant deviation in the CRISPRi screening score are highlighted in the top.

**SFig. 5.**
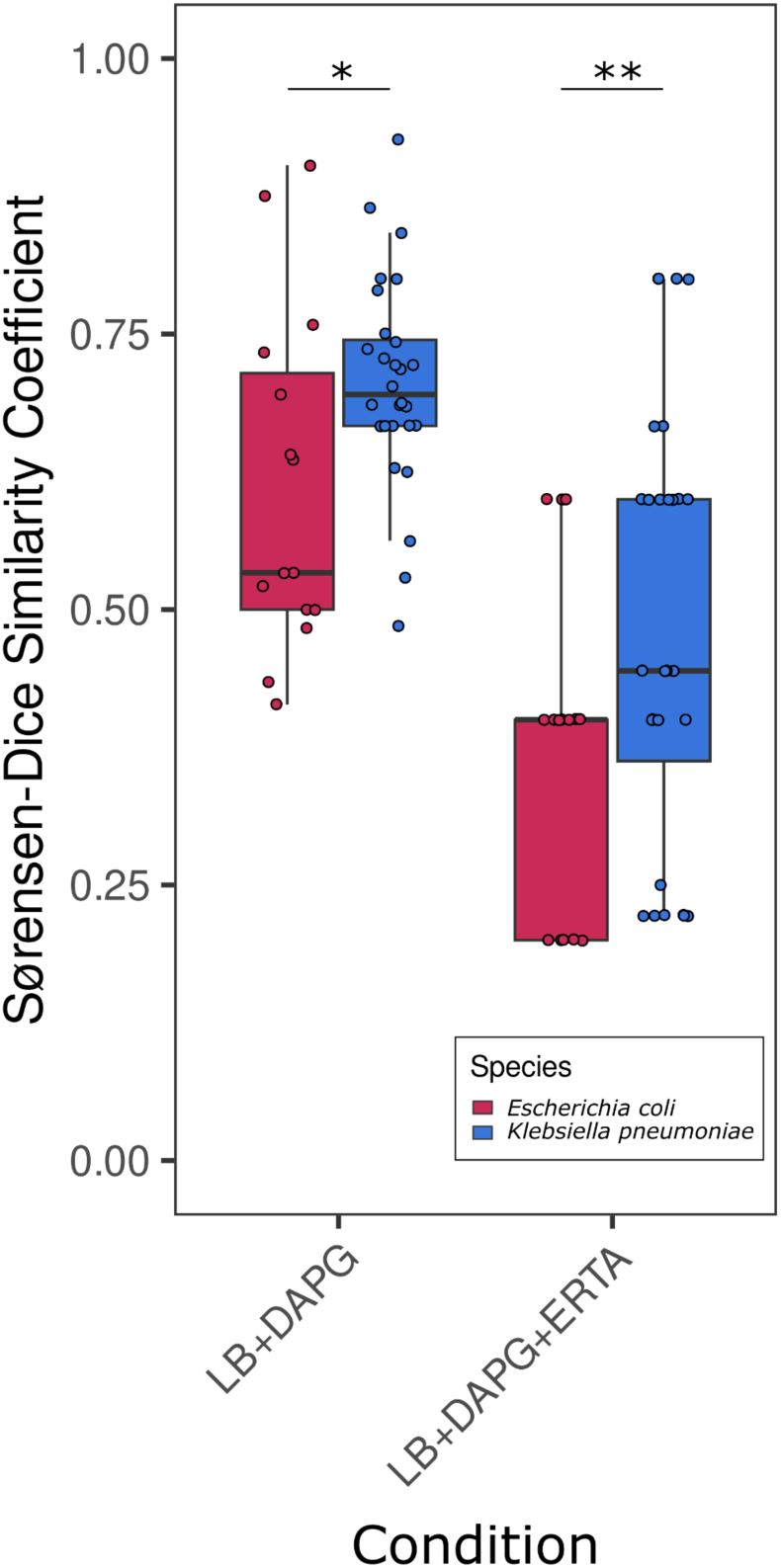
Pairwise similarity between strains (Sørensen-Dice Similarity Coefficient [SDC]). Pairwise comparisons between strains were performed considering significant genes from the CRISPRi screening for each strain either with or without ertapenem (as shown in Fig. 3A but performed individually). SDC=0.5 indicates that 50% of the significant genes between strain X and strain Y coincide. The colors indicate the species. SDC) between strains resulted in moderate (∼0.5) to high (>0.7) values in both species for the CRISPRi screening results in absence of antibiotics. However, in presence of ertapenem, the SDC values obtained were lower, probably due to the lower number of significant targets observed in this condition for each strain (n = 4-5 in ertapenem presence, compared to n = 8-21 in no ertapenem; Suppl. Table X). At least one gene was conserved among all the strains in each species (SDC = 0.2). Asterisks denote significant differences between both species in the different experimental conditions (Welch two sample t-test, t = −2.1694 p-value = 0.04234 without ertapenem; two-sided Kruskal-Wallis test, chi-squared = 5.6326, p-value = 0.01763 with ertapenem). These differences suggest that the response was more conserved in *K. pneumoniae* than in *E. coli*.

**SFig. 6.**
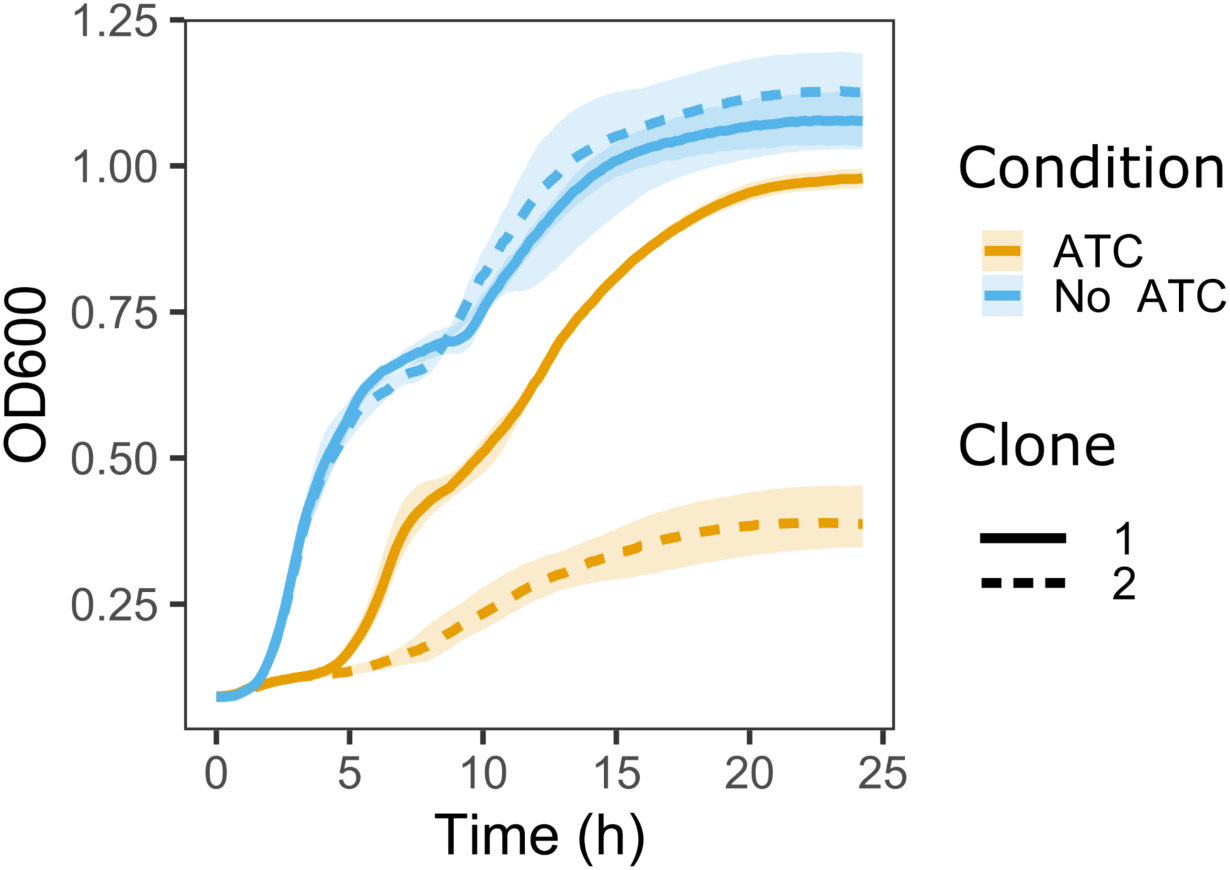
*pemK* expression produces a growth defect in *E. coli* strain EC22. Growth curves of *E. coli* strain EC22 transformed with a vector expressing *pemK* under the control of the pTet promoter (and not carrying pOXA-48). The growth curves are represented as Optical Density (OD) at 600 nm wavelength measured during 24 hours with and without ATC induction. 2 independent clones were assayed per condition, as indicated by the line type (replicates per clone, n = 8). The different behaviours of the clones under ATC induction may be due to escape mutants arising during the experiment.

**SFig. 7.**
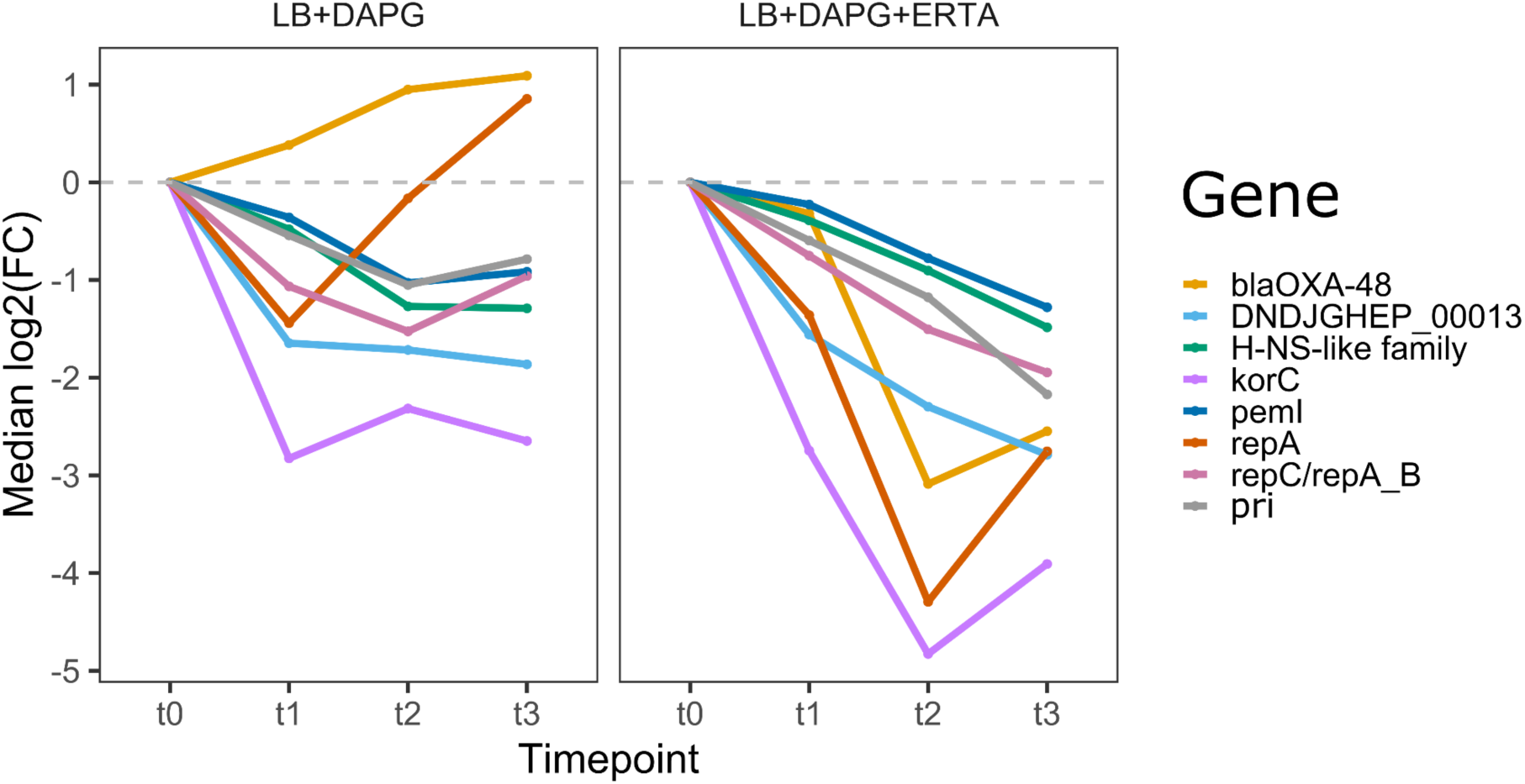
Normalised frequency of guides targeting relevant genes over the screen. The median log2 Fold Change of the guides targeting the most relevant genes from the CRISPRi screen are shown at the different timepoints of the experiment. The values shown correspond to the median value for each gene of the median log2 Fold Change for all the studied strains (n = 14). Each panel represents an experimental condition, either with no antibiotic (left) or with ertapenem (right). The grey dashed line indicates the 0, so that silencing a gene which guide is above this value is beneficial in the given condition (e.g. *bla*_OXA-48_ in absence of antibiotics, orange line in left panel), whereas silencing a gene which guide is below this value is detrimental. The atypical behavior of the *repA* guide in absence of antibiotics can be observed, as it shifts from negative to positive relative frequency in the populations in absence of antibiotics.

**SFig.**
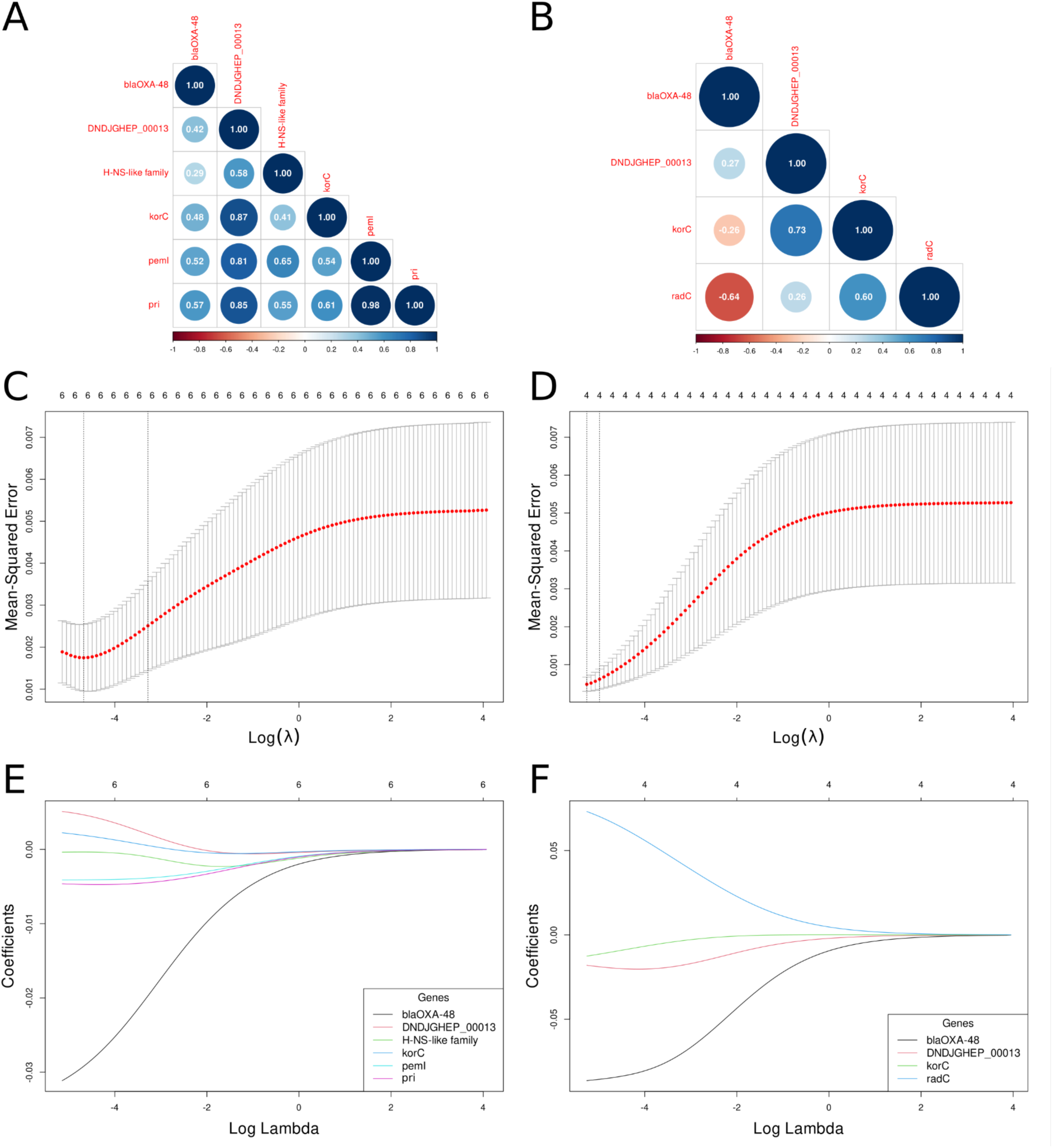
Ridge regression models training parameters. (A,B) Correlation matrices between genes (predictor variables) for *K. pneumoniae* (A) and *E. coli* (B), respectively. Correlation indexes between genes were estimated to check for possible multicollinearity issues in the predictions. Only two genes, *pri* and *pemI*, showed a high correlation index (>0.9), in the *K. pneumoniae* model, which is likely due to the fact that both genes are highly related with plasmid stability. (C,D) Mean-squared error as a function of the lambda parameter. The optimal lambda parameter (which minimises the error) for each model (*K. pneumoniae* [C] and *E. coli* [D]) obtained during the LOOCV is shown). (E, F) Ridge coefficient paths for the genes included in the final Ridge models.

**SFig. 9.**
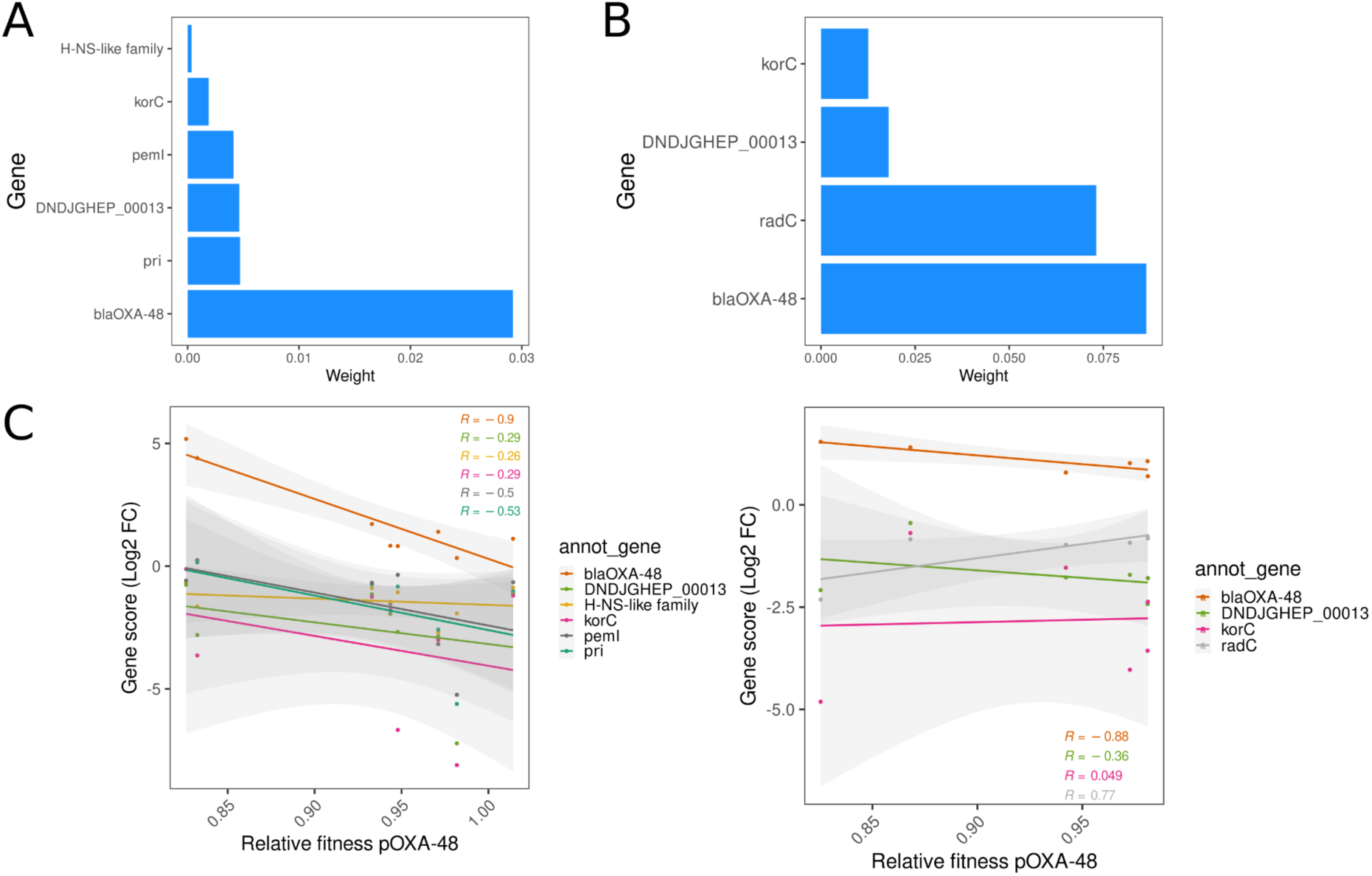
Gene contribution in Ridge Models. (A,B) Weight coefficients for each gene included in the *K. pneumoniae* (A) and *E. coli* (B) Ridge models. For both species, *bla*_OXA-48_ was the gene which most contributed to the relationship between pOXA-48 fitness cost and CRISPRi scores. (C, D) Individual tendencies of the genes included in each Ridge model (C: *K. pneumoniae*, D: *E. coli*).

**SFig. 10.**
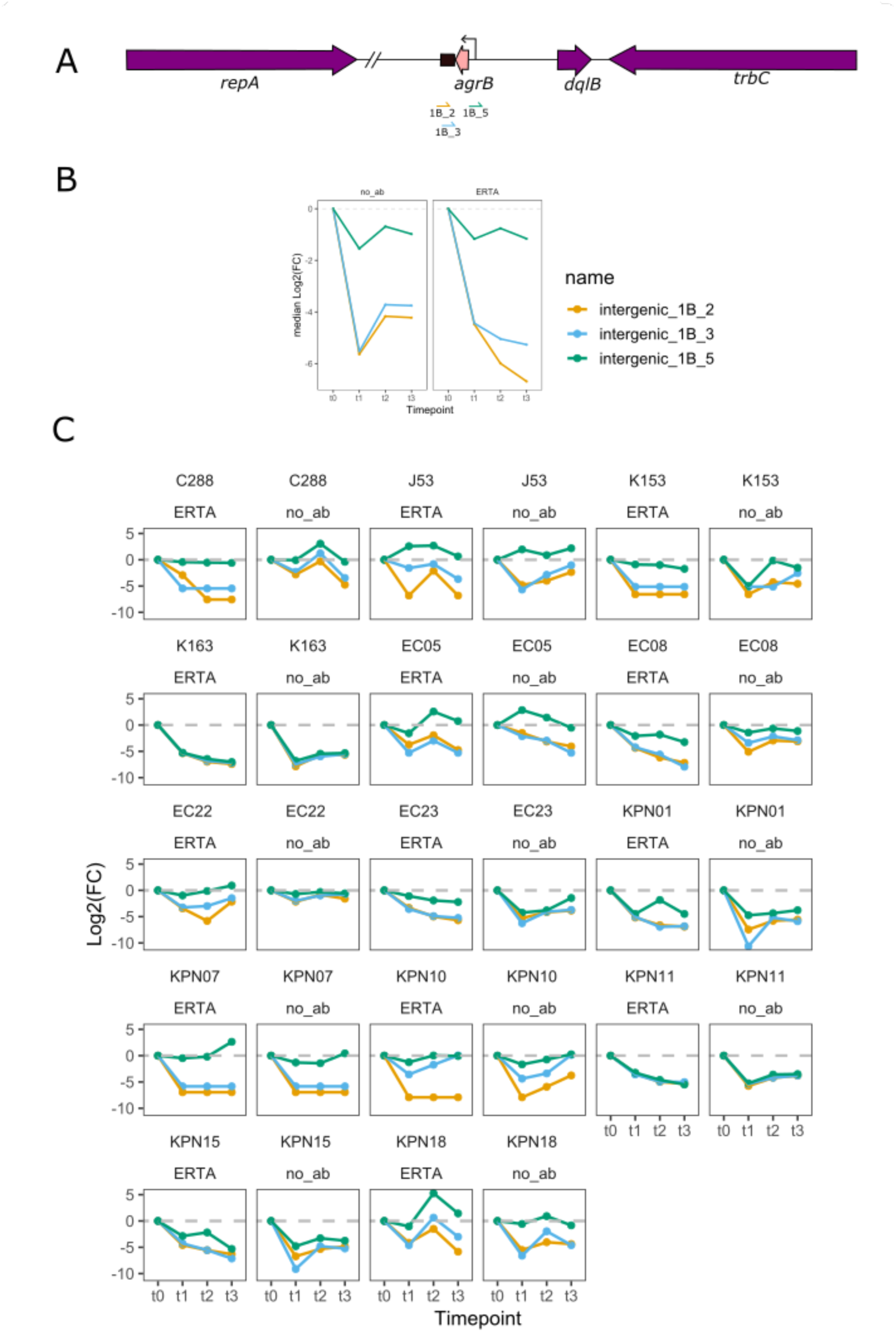
Temporal analysis of log2FC scores for 3 guides targeting *agrB*. Position and log2FC scores of intergenic guides 1B_2, 1B_3 and 1B_5 throughout the CRISPRi experiment. Guide 1B_5 targets the promoter region (indicated with an arrow), and guides 1B_2 and 1B_3 the terminator region (indicated with a black box). (A) Position of the guides in pOXA-48 and annotation of *agrB* described in Baffert *et al.*^37^ (B) Median scores for each guide in all strains of work, in the presence and absence of Ertapenem. (C) Scores for each guide in each individual strain, in the presence and absence of antibiotic.

